# Pre-ciliated tubal epithelial cells are prone to initiation of high-grade serous ovarian carcinoma

**DOI:** 10.1101/2023.12.12.571315

**Authors:** Andrea Flesken-Nikitin, Coulter Q. Ralston, Dah-Jiun Fu, Andrea J. De Micheli, Daryl J. Phuong, Blaine A. Harlan, Amanda P. Armstrong, David McKellar, Sangeeta Ghuwalewala, John C. Schimenti, Benjamin D. Cosgrove, Alexander Yu. Nikitin

## Abstract

The distal region of the uterine (Fallopian) tube is commonly associated with high-grade serous carcinoma (HGSC), the predominant and most aggressive form of ovarian or extra-uterine cancer. Specific cell states and lineage dynamics of the adult tubal epithelium (TE) remain insufficiently understood, hindering efforts to determine the cell of origin for HGSC. Here, we report a comprehensive census of cell types and states of the mouse uterine tube. We show that distal TE cells expressing the stem/progenitor cell marker *Slc1a3* can differentiate into both secretory (*Ovgp1*+) and ciliated (*Fam183b*+) cells. Inactivation of *Trp53* and *Rb1*, whose pathways are commonly altered in HGSC, leads to elimination of targeted *Slc1a3*+ cells by apoptosis, thereby preventing their malignant transformation. In contrast, pre-ciliated cells (*Krt5*+, *Prom1+*, *Trp73*+) remain cancer-prone and give rise to serous tubal intraepithelial carcinomas and overt HGSC. These findings identify transitional pre-ciliated cells as a previously unrecognized cancer-prone cell state and point to pre-ciliation mechanisms as novel diagnostic and therapeutic targets.

## Background

Ovarian cancer is the fifth leading cause of death for women in the United States ^1^. High-grade serous carcinoma (HGSC) is the most common and aggressive type of ovarian cancer ^2,3^. Over 80% of HGSC are detected at advanced stage and have limited treatment options ^2,4,5^. This is attributed to latent progression of the disease with lack of early symptoms and detection methods ^2^. Detection and treatment of HGSC at earlier stages could be crucial to improving the prognosis of patients with this malignancy. However, identification of new diagnostic markers and therapeutic targets is hindered by our inadequate knowledge about the cells in which HGSC originates and the mechanisms underlying disease initiation.

The location of HGSC initiation has long been debated, but the emerging consensus is that both the ovarian surface epithelium (OSE) and the tubal epithelium (TE) of the uterine tube, also known as the oviduct or Fallopian tube, have potential to progress into HGSC ^6–11^. While cancer-prone stem/progenitor cells have been described for the OSE ^12,13^, the cell of origin of HGSC arising from TE remains unclear. It has been shown that the majority of familial HGSC cases may begin with the appearance of serous tubal intraepithelial carcinomas (STIC)^6^. These lesions are found exclusively in the distal region of the uterine tube ^2,14,15^. STICs lack ciliation, express the transcriptional factor PAX8, and harbor mutations in the *TP53* gene (also known as *Trp53* in the mouse), which encodes for p53. *TP53* mutations are the most frequent genetic alterations in HGSC, being present in over 96% of cases ^16,17^. Consistent with these observations, STICs can be induced by inactivation of tumor suppressor genes commonly associated with human HGSC, such as *Trp53*, *Brca1*, *Brca2* and *Rb1* in *Pax8*-expressing tubal epithelial cells of the mouse uterine tube ^18^.

In both humans and rodents, uterine tubes consist of distal (infundibulum, ampulla and ampullary-isthmic junction) and proximal regions (isthmus and intramuscular utero-tubal junction). The uterine tube is formed by the simple pseudostratified TE surrounded by a thin stromal layer, two circular smooth muscle layers and the mesothelium. Two main cellular components of the TE are ciliated cells (also known as multi-ciliated cells), predominantly located in the distal regions of the tube, and secretory cells, which are more abundant in the proximal regions. Additionally, there are basal cells, representing intraepithelial T-lymphocytes and peg cells. Peg cells have been described as either exhausted secretory cells or CD44+ progenitor cells ^19–21^.

Previous mouse lineage-tracing studies have reported that cells expressing *Pax8* have the capacity to self-renew and differentiate into ciliated cells in both distal and proximal regions after labeling during embryonic or prepubertal development ^22^. However, recent studies based on immunophenotyping, lineage tracing, and limited single cell RNA-sequencing (scRNA-seq) suggest presence of distinct cell lineages in the distal (*Pax8*+) and the proximal (*Pax2*+) regions of the adult mouse uterine tube ^23^.

It is well established that many types of cancer arise from stem cell niches ^24–26^. Previously, using a genetically defined mouse model, we have shown that ovarian surface epithelium (OSE) stem/progenitor cells can be efficiently transformed after inactivation of tumor suppressor genes *Trp53* and *Rb1* and lead to HGSC formation ^12,13,27^. Notably, tumors arising from non-stem OSE cells were slow-growing and non-metastatic. However, in other cancer types, neoplasms may originate from differentiated or transitional state cells that have acquired some stem cell properties ^27–29^. It has been hypothesized that some ovarian carcinomas may arise from the ciliated cell lineage ^30^. However, no direct experimental data has been offered to support this idea.

Here, we conducted scRNA-seq to establish a comprehensive census of cell types found in the mouse uterine tube. Proximal and distal sections were sequenced separately to investigate characteristics underscoring the distal region’s predisposition towards cancer initiation. By using a combination of computational lineage trajectory projections and genetic cell fate, we identified a TE stem/progenitor cell population and interrogated unique epithelial cell states for their propensity for malignant transformation. These studies revealed that pre-ciliated cells may serve as a specific cell state susceptible to malignant transformation.

## Results

### Census of cell types of the mouse uterine tube

The mouse uterine tube is divided along the distal and proximal axis (Figure 1a). We collected 62 uterine tubes from 31 adult mice, separated distal and proximal regions, and processed them for scRNA-seq (Supplementary Figures 1-3) by region. After sequencing and data preprocessing of distal region samples, 16,583 high quality cells remained. Following Harmony integration^31^, we identified 18 clusters by Louvain clustering contributing to epithelial, stromal, and immune compartments, which we visualized using the uniform manifold approximation and projection (UMAP) (Figure 1b). Features defining each cluster were found using general markers known to define each subtype (Figure 1c). Epithelial cells were characterized by epithelial markers (*Epcam* and *Krt8*), and further epithelial specificity was achieved with secretory (*Ovgp1*) and ciliated (*Foxj1*) markers. Additional cell types were noted within our dataset to be mesothelial and luteal cells.

**Figure 1.**
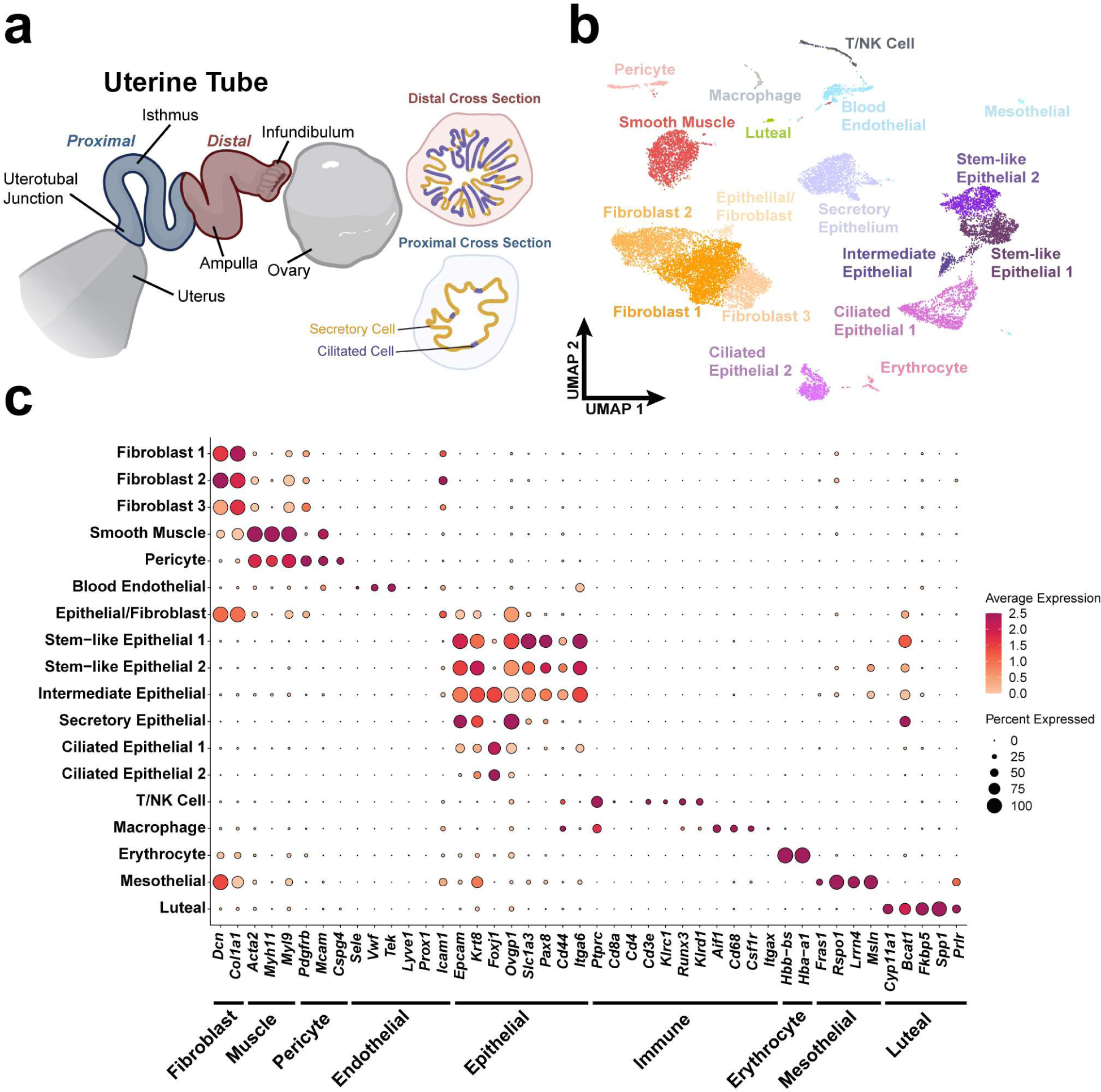
Census of cell types of the mouse uterine tube. (**a**) Diagram of a partially uncoiled mouse uterine tube. The proximal region contains few ciliated cells and extends from the intratubal junction to the ampulla. The distal region consists of the ampulla and the infundibulum where ciliated cells are abundant. (**b**) UMAP visualization of the cell types identified in a pool of 16,583 distal cells from which high quality sequence data was obtained. Visualization of a UMAP embedding (**c**) Dot plot representation of genes associated with various tissue types to validate cell type identification.

### Characterization of distal epithelial cell states

To further investigate the unique cell states of the tubal epithelium, we subset the epithelial clusters of the uterine tube cell types. After processing the epithelial subset, we identified 9 clusters consisting of ciliated, secretory, stem-like, and transitional cell states (Figure 2a and Supplementary Figure 3a-c). General epithelial markers (*Epcam*, *Krt8*), secretory markers (*Ovgp1*, *Sox17*), and ciliated markers (*Fam183b, Foxj1*) were used to classify groups of epithelial cells that are known to reside in the tubal epithelium (Figure 2b). Additional features were found using the Wilcoxon Rank Sum Test to determine cluster-specific markers. In agreement with previous studies ^20,32^, ciliated cells and their precursors were mainly located in the distal region (32% distal vs 10% proximal), while secretory cells predominantly populated the proximal region (30% distal vs 65% proximal; Supplementary Table 1). Correlation between the clusters identified in proximal and distal regions only showed strong correlation between a subset of secretory cell states between the two regions (Supplementary Figure 3d). In the distal region *Pax8* expression was mainly detected in stem/progenitor cell cluster (81% of cells) and transitional pre-ciliogenic state cells (49%). Only 3% of ciliated cells and 28% of secretory cells were *Pax8+* (Figure 2b, Supplementary Figure 4 and Supplementary Table 1). In the proximal region 67% of putative stem/progenitor cells expressed *Pax8*, while both secretory and ciliated clusters contained about 29% of *Pax8+* cells each.

**Figure 2.**
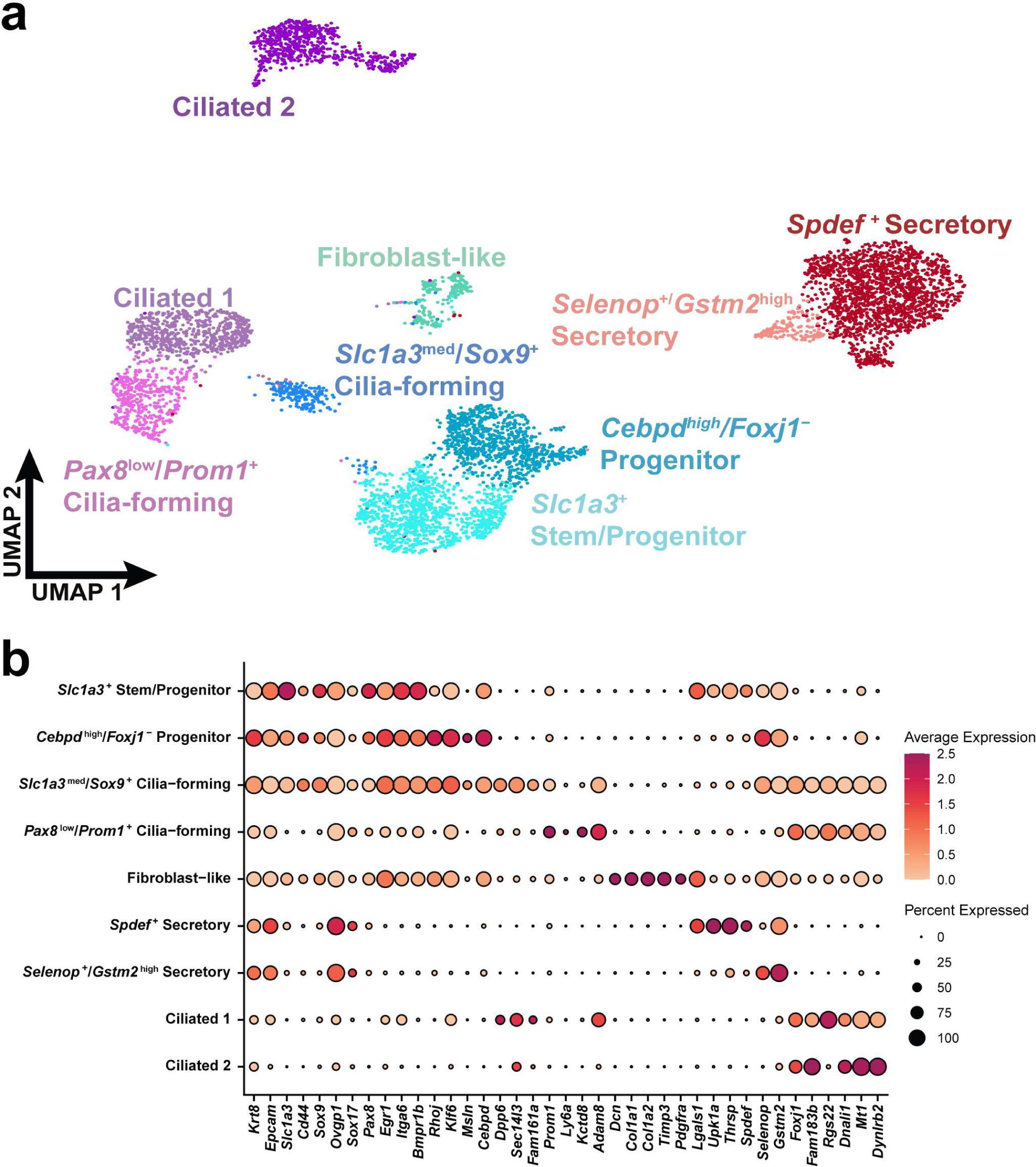
Characterization of distal epithelial cell states. (**a**) Epithelial cells were identified by their *Epcam* and *Krt8* expression and the subset was represented within the UMAP embedding. (**b**) Dot plot reflecting highly expressed, specific markers of each identified epithelial cell cluster.

General expression of stem-like markers indicated that *Slc1a3* might serve as a potential marker for epithelial stem/progenitor cells that gives rise to both secretory and ciliated cells (Figure 2b and Supplementary Figure 4). *Slc1a3* was previously described as a marker of cell population containing distinct skin epithelial stem cells ^33,34^. We employed PHATE (Potential of heat diffusion for affinity-based transition embedding) ^35^ to better visualize how tubal epithelial cells progress in a two-dimensional representation similar to UMAP (Figure 3a). A pseudotime trajectory was calculated with Monocle3 ^36^ and overlayed on the PHATE embedding (Figure 3b). In the PHATE representation, *Slc1a3* expression is located at the center with secretory and ciliated branches split by the suspected stem/progenitor cell state (Figure 3c). *Pax8* expression is present in *Slc1a3*+ cells but also extends towards early cilia-forming cells (Figure 3d).

**Figure 3.**
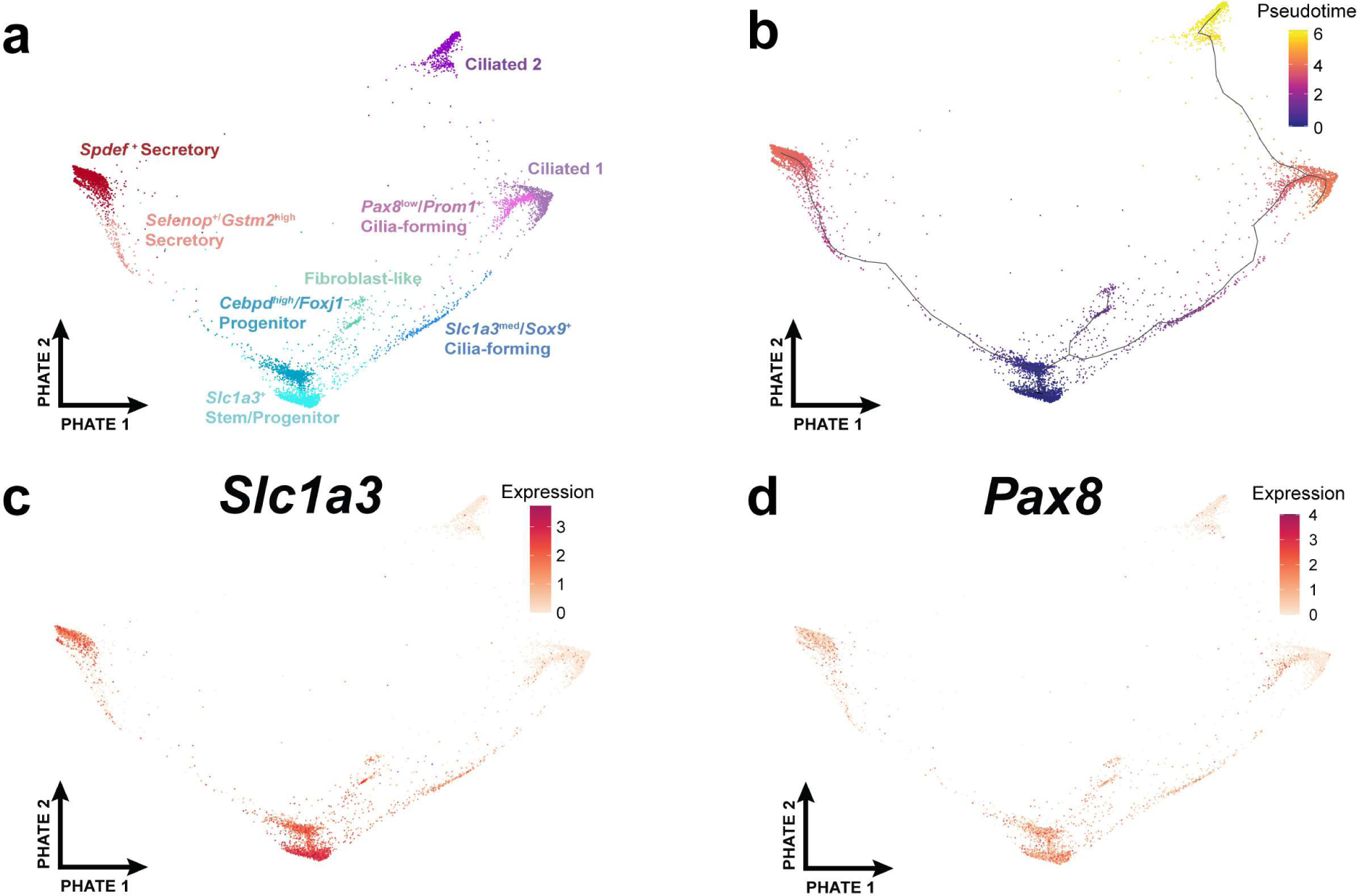
Feature expression over the PHATE embedding. (**a**) A differentiation trajectory among epithelial cells visualized through the PHATE dimensional reduction technique. (**b**) Monocle3 pseudotime analyses calculated over the PHATE embedding. (**c** and **d**) The expression of *Slc1a3* (c) and *Pax8* (d) visualized over the PHATE embedding.

### *Slc1a3*+ epithelial cells are stem/progenitor cells for the TE

To test if *Slc1a3*+ epithelial cells are stem/progenitor cells of the TE, cell lineage-tracing and organoid studies were conducted in Slc1a3-CreERT Ai9 mice. In these mice, administration of tamoxifen allows for Cre-*loxP* mediated induction of tdTomato expression in *Slc1a3*+ cells (Figure 4a). One day after tamoxifen induction tdTomato expression was mainly detected in the distal TE (Figure 4b and c). Within 30 days after tamoxifen injection, tdTomato+ cells expanded to form clusters and persisted for over one year (Figure 4b and d). These cells showed expression of secretory (OVGP1) and ciliated (FAM183B) cell markers (Figure 4b, e and f).

**Figure 4.**
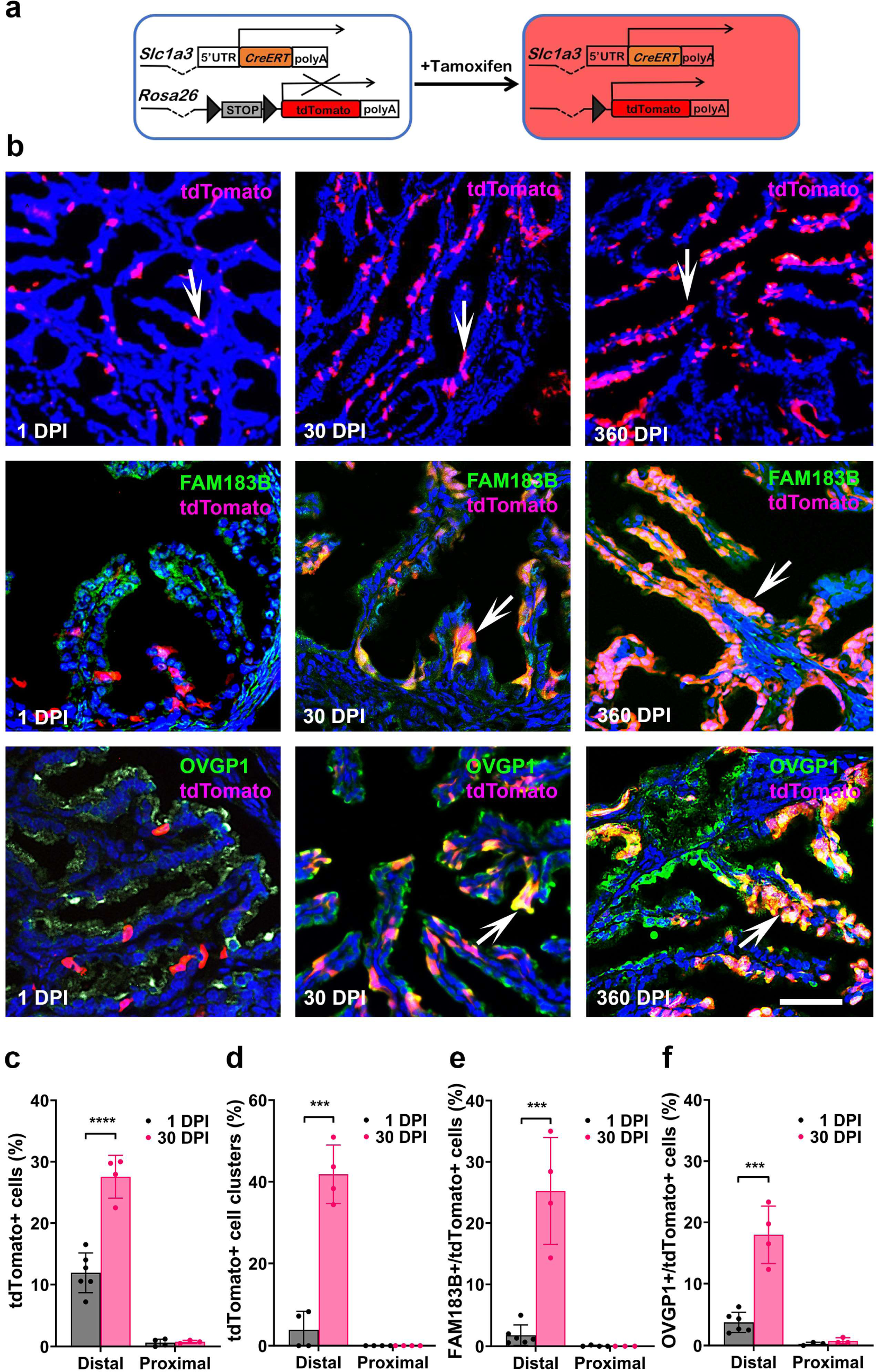
Lineage tracing of *Slc1a3+* stem/progenitor cells. (**a**) Experimental design. Mice containing Slc1a3-CreERT and Ai9 reporter are injected with tamoxifen to induce expression of modified red fluorescent protein (tdTomato) in cells expression *Slc1a3*. (**b**) Top row, tdTomato expression (red, arrows) 1, 30 and 360 days post induction (DPI) with tamoxifen. tdTomato+ progeny of *Slc1a3*+ cells express differentiation markers for ciliated (FAM183B, middle row, arrows) and secretory (OVGP1, bottom row, arrows) cells. Counterstaining with DAPI. Scale bar represents 100 μm (top row), and 50 μm (middle and bottom row). (**c**) Quantification of cells expressing tdTomato in the distal and proximal regions of the uterine tube 1 and 30 DPI. (**d**) Frequency of clusters containing at least 3 adjacent tdTomato+ cells. (**e** and **f**) Quantification of cells expressing tdTomato with either ciliated (FAM183B) or secretory (OVGP1) cells 1 and 30 DPI. ***P < 0.001, ****P < 0.0001, two-tailed unpaired t-tests. All error bars denote s.d. n=3 or more in each group.

**Figure 5.**
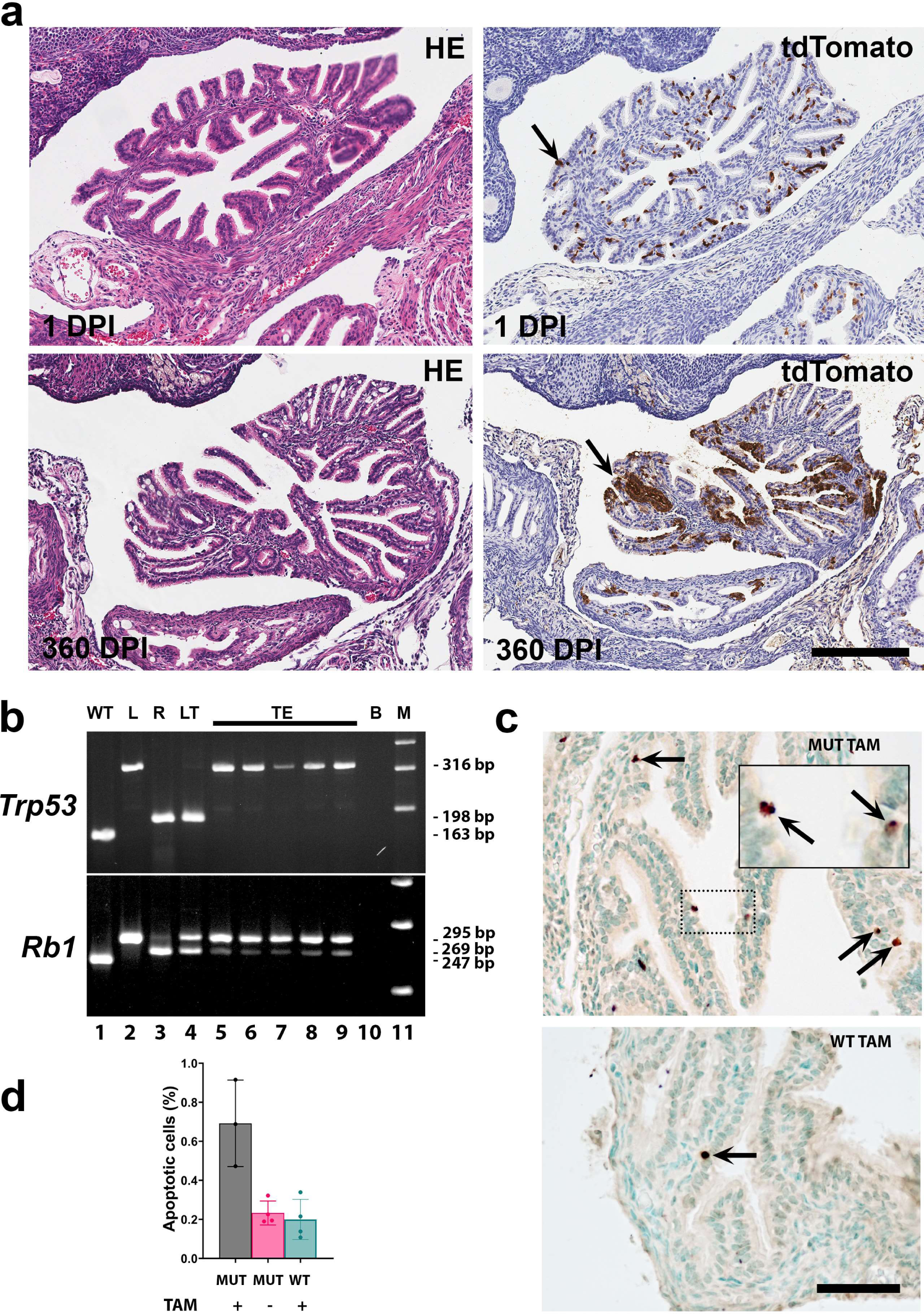
*Slc1a3+* epithelial cells are not cancer-prone. (**a**) tdTomato+ tubal epithelial cells in mice containing Slc1a3-CreERT with floxed *Trp53* and *Rb1* genes, and an Ai9 reporter 1 and 360 days post induction (DPI) with tamoxifen. Hematoxylin and Eosin (HE) staining (left column) and immunostaining for tdTomato (arrows, brown color), Elite ABC method, hematoxylin counterstaining. (right column). Bar, all images 200 μm. (**b**) PCR analysis of *Trp53* and *Rb1* gene structure in the same samples of microdissected cells from the tubal epithelium (TE, lanes 5-9) and lung neoplasm (LT, lane 4) of Slc1a3-CreERT *Trp53^loxP/loxP^ Rb1^loxP/loxP^* Ai9 mice collected one year after tamoxifen induction. Samples with known gene structure (wild-type, WT, lane 1, floxed gene, L, lane 2, and recombinant gene in gastric cancer of *Lgr5*^eGFP−Ires−CreERT2^*Trp53^loxP/loxP^Rb1^loxP/loxP^*Ai9 mice^27^, R, lane 3). 316-, 198-, and 163-bp fragments are diagnostic for floxed, excised, and wild-type alleles of the *Trp53* gene, respectively. 295-, 269, and 247-bp fragments are diagnostic for floxed, excised, and wild-type alleles of the *Rb1* gene, respectively. B, blank control (lane 10), M, DNA marker (Lane 11), (**c)** Apoptotic cells (arrows) in the tubal epithelium one day after tamoxifen induction of Cre-mediated inactivation of *Trp53* and *Rb1* in Slc1a3-CreERT *Trp53^loxP/loxP^ Rb1^loxP/loxP^* Ai9 mice (MUT TAM) and littermates without Slc1a3-CreERT (WT TAM). Dot-line rectangle indicates respective location of cells shown in the inset in the top image. TUNEL assay, methyl green counterstaining. Bar, 50 μm and 21 µm, inset. (**d**) Quantification of apoptotic cells one day after administration of tamoxifen (TAM+, n≥3) or vehicle (TAM-, n=4) in Slc1a3-CreERT *Trp53^loxP/loxP^ Rb1^loxP/loxP^* Ai9 mice (MUT) or littermates without Slc1a3-CreERT (WT). (d) One way ANOVA P=0.0024, Error bars denote s.d.

Consistent with previous studies of mouse and human TE ^19,37,38^, organoids were preferentially formed by mouse distal TE (Supplementary Figure 5a-d). Furthermore, the organoid forming ability of tdTomato+ cells was significantly higher than tdTomato-cells isolated from the distal region (Supplementary Figure 5e-g).

In sum, based on general expression of stem-like markers, computational lineage trajectory projections, cell lineage tracing, and organoid assays, *Slc1a3*+ epithelial cells represent stem/progenitor cells for the distal TE, and contribute to the long-term maintenance of both ciliated and secretory epithelial cells within the uterine tube.

### *Slc1a3*+ epithelial cells are not cancer-prone

To investigate the role of *Slc1a3*+ cells in malignant transformation we have prepared Slc1a3-CreERT *Trp53^loxP/loxP^ Rb1^loxP/loxP^* Ai9 mice. However, no TE neoplastic lesions were observed in these mice (n=19) by one year after tamoxifen administration (Figure 4a). Large clusters of TE cells expressed tdTomato at the time of collection. Cre-mediated excision of *Rb1* but not *Trp53* was detected by microdissection-PCR assay in all tested samples (Figure 4b). Evaluation of TE has shown increased apoptotic rate one day after tamoxifen administration (Figure 4c and d). Thus, *Trp53* inactivation leads to elimination of targeted *Slc1a3*+ cells by apoptosis, thereby explaining their resistance to malignant transformation.

### Pre-ciliated cells are susceptible to malignant transformation

Consistent with previous reports ^18,39^, we have observed that TE can be transformed by conditional inactivation of *Trp53* and *Rb1* in cells expressing *Pax8* (Figure 6a and b, Supplementary Table 2). STICs (6 out of 12 cases) and/or HGSC (3 out of 12) were detected in 58% of mice by 400 days post induction (DPI). These data suggest that a subset of *Pax8*+ cells is susceptible to malignant transformations, distinct from *Slc1a3*+ stem/progenitor cells that do not give rise to malignancies.

**Figure 6.**
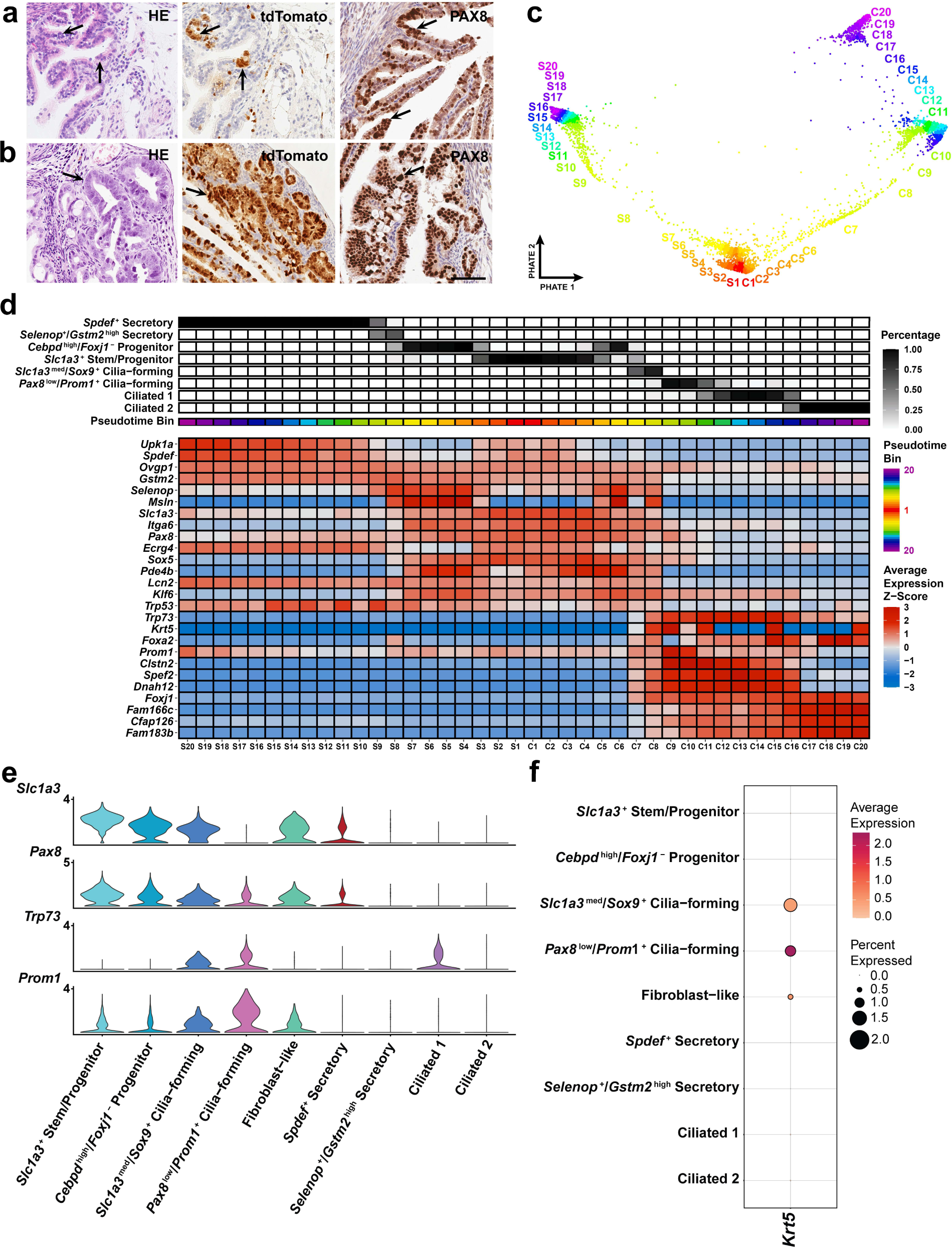
Identification of cancer-prone cell states. (**a** and **b**) Early (a) and advanced (b) neoplastic lesions (arrows) in Pax8-rtTA Tre-Cre *Trp53^loxP/loxP^ Rb1^loxP/loxP^* Ai9 mice. Hematoxylin and Eosin (HE) staining (left column) and immunostaining for tdTomato (middle column, brown color), and PAX8 (right column, brown color). Elite ABC method, hematoxylin counterstaining. Bar, 60 µm. (**c**) Pseudotime binning along the PHATE embedding to visualize how bins are assigned. (**d**) Inferred pseudotime trajectories of secretory and ciliated epithelial cell lineages. The lineages extend from S1 and C1 to S20 and C20 respectively, where S20 and C20 are presumed to be a more differentiated cell state. The percent abundance of each cell type contributing to each pseudotime bin is reflected in black. The average z-scored expression was calculated for each gene to identify genes that best represent smaller transitional states within each lineage. Each pseudotime bin is equally sized and consists of about 150 cells. (**e**) Log-normalized expression of *Slc1a3, Pax8, Trp53 and Prom1* visualized in a violin plot of epithelial cell clusters. (**f)** Dot plot of *Krt5* expression among epithelial cell populations.

To define a unique population of *Pax8*-expressing cells implicated in malignant transformation, we leveraged our scRNA-seq analysis to identify a group of cells expressing *Pax8* but not *Slc1a3* in the distal region of the uterine tube. Strikingly, the only cell state exhibiting *Pax8* expression without concomitant *Slc1a3* expression displayed features consistent with pre-ciliated cells (Figure 6c-e). We subset the branched trajectory of secretory and ciliated cells from *Slc1a3*+ progenitors and formed 20 pseudotime bins equally divided along the trajectory to total about 150 cells per bin (Figure 6c-d). We have found that *Pax8+*, *Slc1a3*-cells are present as a transitional state along the ciliogenic lineage. Such cells are characterized by expression of *Krt5, Prom1* and *Trp73* (Figure 6d-f). Notably, expression of these genes negatively correlates with *Trp53* expression, thereby suggesting reduced requirement for *Trp53* during ciliogenesis (Figure 6d and Supplementary Figure 6). At the same time, the majority of other putative driver genes identified in human HGSC^13,40^, including *Rb1*, *Nf1*, and *Pten*, are preferentially expressed in stem/progenitor populations along the pre-ciliogenesis trajectory. Expression of *Brca1, Brca2* and *Csmd3* is mainly present in cilia-forming cells (Supplementary Figure 6).

To test if pre-ciliated cells are susceptible to malignant transformation, we prepared Krt5-CreERT *Trp53^loxP/loxP^ Rb1^loxP/loxP^* Ai9 mice. By 200 days after induction (DPI) 70% of mice developed STIC (10 out of 16) and/or HGSC (3 out of 16; Fig. 7a-d and Supplementary Table 2). Notably, 7 out of 10 STIC lesions (70%) were located near TE-mesothelial junctions, similar to previously described location of human STICs in close vicinity to the tubal-peritoneal junction^14,15^. The earliest atypical lesions were observed at 104 DPI as compared to 154 DPI in experiments with inactivation of *Trp53* and *Rb1* in *Pax*8+ cells. Notably, according to tdTomato cell labeling at 1 DPI, the pool of targeted *Krt5*+ cells (0.83%) was significantly smaller than that of *Pax8*+ cells (P=0.0007; Figure 7d and Supplementary Table 2). These observations support the notion that cells in a transitional state along ciliogenesis are cancer-prone.

**Figure 7.**
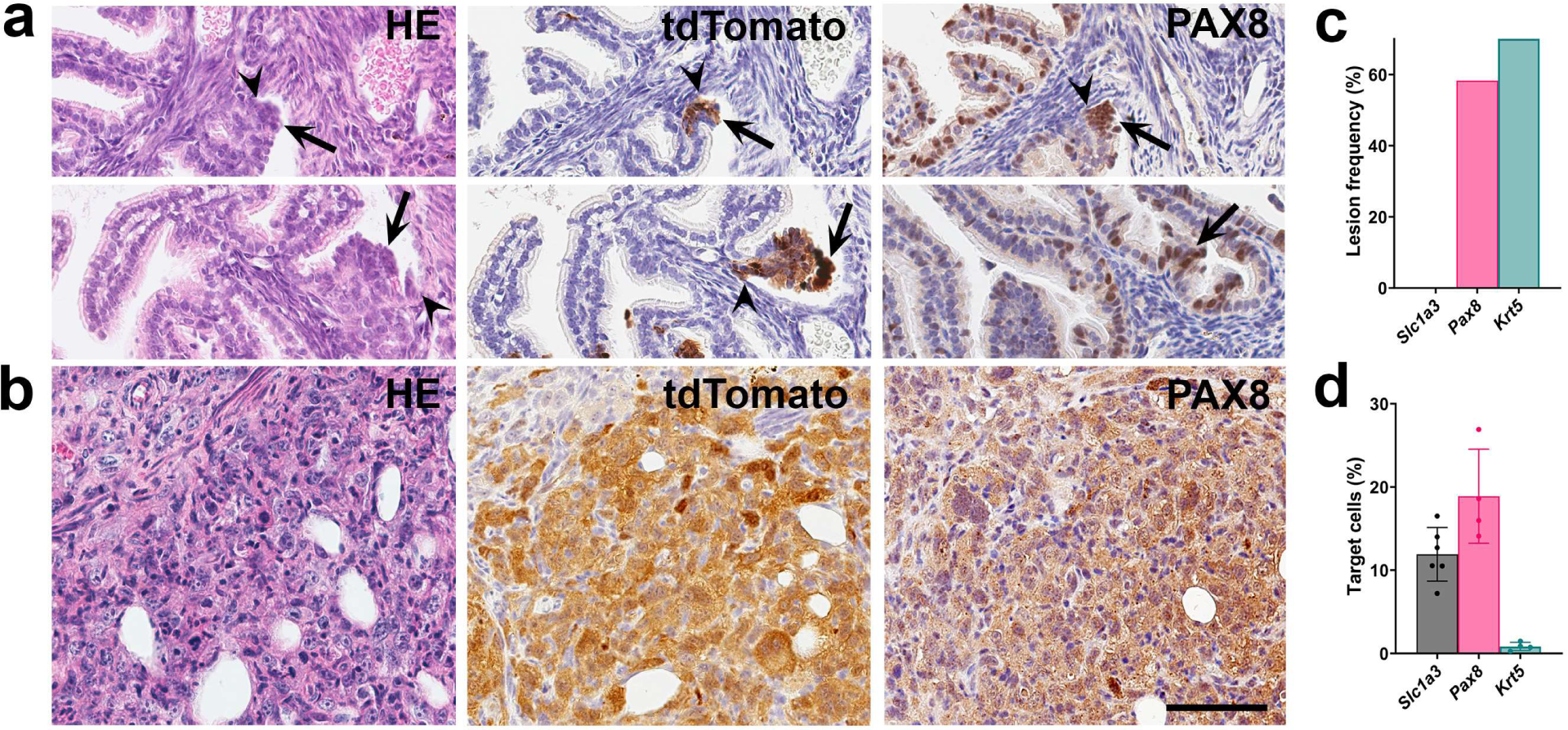
Transitional pre-ciliated *Krt5*+ cells are highly susceptible to malignant transformation. (**a** and **b**) Early (a, arrows) and advanced (b) neoplastic lesions in Krt5-CreERT *Trp53^loxP/loxP^ Rb1^loxP/loxP^* Ai9 mice. Arrowheads, TE-mesothelial junctions. Hematoxylin and Eosin (HE) staining (left column) and immunostaining for tdTomato (middle column, brown color), and PAX8 (right column, brown color), Elite ABC method, hematoxylin counterstaining. Bar, 60 µm. (**c** and **d**) Frequency of neoplastic lesions (c) and target cells (d) in Slc1a3-CreERT (Slc1a3), Pax8-rtTA Tre-Cre (Pax8) and Krt5-CreERT (Krt5) mice carrying floxed *Trp53* and *Rb1* genes. d, ANOVA with Tukey-Kramer Multiple Comparisons P value < 0.0001 ** Error bar denotes s.d.

## Discussion

Our single-cell transcriptomics allowed for unique cell states and cell lineage hierarchy to be identified in the context of healthy adult mouse uterine tubes. Further experimentation has confirmed that *Slc1a3*+ cells give rise to both secretory (OVGP1+) and ciliated (FAM183B+) cells in the distal region of the uterine tube. Notably, expression of *Pax8* was mainly detected in stem/progenitor cells and transitional cell states. These findings confirm previous observations that *Pax8*+ cells can differentiate into ciliated cells ^22^. However, they also indicate that mature secretory cells do not give rise to ciliated cells.

We could not observe any significant contribution of *Slc1a3*+ cells to cells of the proximal uterine tube. This can be explained by the existence of developmentally distinct cell lineages forming the proximal region ^23^.

Our current study shows that unlike stem/progenitor cells of the ovarian surface epithelium, *Slc1a3+* TE stem/progenitor cells do not undergo malignant transformation after Cre-mediated inactivation of *Trp53* and *Rb1*. This was not due to the lack of Cre activity in *Slc1a3+* cells. Long-term contribution of tdTomato+ cells to the tubal epithelium was observed in all uterine tubes by 360 days after induction of Cre expression. Furthermore, all samples contained an excised *Rb1* allele. However, we observed only intact floxed *Trp53* in tdTomato+ TE cells and increased apoptotic index one day after induction of Cre expression, supporting the notion that inactivation of *Trp53* is incompatible with long-term survival *Slc1a3+* cells.

By comparative evaluation of *Pax8+* cells lacking concomitant *Slc1a3* expression, we identified a cancer-prone cell population along the ciliogenic lineage. This observation is consistent with STIC prevalence in the distal, ciliated cell-rich, region of the uterine tube ^2,14,15^. Furthermore, the majority of HGSC putative driver genes ^13,40^, either preferentially expressed in stem/progenitor cells with a bias towards pre-ciliogenic trajectory (*Rb1, Nf1* and *Pten)*, or mainly detected in cilia-forming cells (*Brca1*, *Brca2* and *Csmd3*). Thus, aberrations in these genes may have the most transformation potential in the context of pre-ciliogenic cell state, as opposed to secretory differentiation.

Our previous studies have shown that ciliated *Foxj1*+ cells are not transformed by inactivation of *Trp53* and *Rb1* ^39^. Hence cancer-susceptibility of TE cells deficient for these genes is limited to a transitional pre-ciliated cell state marked by expression of *Krt5, Prom1* and *Trp73.* All three genes have been linked to human ovarian cancer. Expression of both *KRT5* and *PROM1* are associated with HGSC progression ^41,42^. *TP73,* encoding human p73, is upregulated in many cases of epithelial ovarian cancers and modulates the sensitivity of ovarian cancers to chemotherapy (reviewed in ^30^).

Notably, *Trp73* is a critical regulator of ciliogenesis and its inactivation results in ciliopathies ^43–45^. Among downstream targets of *Trp73,* are micro-RNAs of the miR-34 family ^46^. These micro-RNAs are also an essential component for ciliogenesis ^47^. At the same time, inactivation of *mir-34* family of genes is commonly observed in HGSC ^48^, suggesting a link between regulation of ciliogenesis and ovarian cancer.

p73 encoded by *TP73/Trp73* is a homolog of the tumor suppressor transcriptional factor p53. Some functions of p53 and p73 are redundant, as evident by the ability of p73 to activate p53-regulated genes in growth suppression and apoptosis induction. However, other functions do remain unique to either p53 or p73, as evidenced by the lack of ciliation abnormalities in mice null for *Trp53* and rare mutations of *TP73* in cancers ^49^. Consistent with distinct functions of p53 and p73 in TE cells, we have observed that *Trp73* expression negatively correlates with *Trp53* expression.

Our studies show that *Slc1a3+* stem/progenitor cells express *Trp53* and its inactivation is incompatible with their survival. Unlike *Slc1a3*+ cells, transitional pre-ciliated cells express both *Trp53* and *Trp73*. This suggests that *Trp73* expression protects cells from apoptosis but not from genomic instability triggered by *Trp53*, thereby leading to the preferential transformation of pre-ciliated cells. Further studies are required to understand how transition from p53-regulated programs in stem/progenitor cells to p73 controlled ciliogenesis programs may predispose to cancer.

We have observed preferential STICs location near the junction between TE and mesothelium, in close similarity to human STICs^14,15^. Junction areas (aka epithelial transitional zones) are anatomically defined regions of organs where two different types of epithelial tissue meet. Such junction areas are known to be predisposed to cancer in many locations such as the ovarian hilum region, the gastric squamo-columnar junction, the corneal limbus region, the anal canal, and the uterine cervix. They also contain stem cell niches reviewed in ^25^. The existence of a cancer-prone cells in junction sites has been demonstrated definitively in the ovarian hilum region and the gastric squamo-columnar junction ^12,13,27^. Our current observation reinforces the notion that high cancer frequency at the junction areas can be explained by the presence of cancer-prone stem cell niches. Further studies should specifically evaluate the mechanisms facilitating preferential transformation of pre-ciliogenic cells at the tubal-mesothelial junction.

In sum, our study establishes the cell hierarchy, cell lineage dynamics, and identity of stem/progenitor cells in the distal tubal epithelium. Furthermore, we show resistance of TE stem/progenitor cells to malignant transformation and provide direct experimental evidence for cancer propensity of cells in the transitional pre-ciliated state. These findings explain the preferential appearance of neoplastic lesions in the distal region of the uterine tube and point to the pre-ciliation state as a novel diagnostic and therapeutic target.

## Methods

### Experimental animals

The Tg(Slc1a3-cre/ERT)1Nat/J (Slc1a3-CreERT, Stock number 012586), Tg(Pax8-rtTA2S*M2)1Koes/J (Pax8-rtTA, stock number 007176), Tg(tetO-Cre)1Jaw/J (Tre-Cre, stock number 006234), Tg(KRT5-cre/ERT2)2Ipc/Jeldj (K5-Cre-ERT2, Stock number 018394), *Gt(ROSA)26Sor^tm9(CAG–tdTomato)Hze^* (*Rosa-loxp-stop-loxp-*tdTomato/Ai9, Stock number 007909), and C57BL6 (Stock number 000664) mice were obtained from The Jackson Laboratory (Bar Harbor, ME, USA). The *Trp53^loxP/loxP^* and *Rb1^loxP/loxP^* mice, which have *Trp53* and *Rb1* genes flanked by *loxP* alleles, respectively, were a gift from Dr. Anton Berns. All the experiments and maintenance of the mice were following ethical regulations for animal testing and research. They were approved by the Cornell University Institutional Laboratory Animal Use and Care Committee.

### Doxycycline and tamoxifen induction

For estimation of target cell frequency 6 to 8 week-old Pax8-rtTA Tre-Cre Ai9 mice received a single dose (12 μl g^-1^ body weight) of doxycycline (6.7 mg ml^-1^ in sterile PBS) by intraperitoneal (i.p.) injection. Identical injection schedule was used for tumor induction experiments with 6 to 10 week-old Pax8-rtTA Tre-Cre *Trp53^loxP/loxP^ Rb1^loxP/loxP^* Ai9 mice and control mice. In our cohort, about 10% of TRE-Cre *Trp53^loxP/loxP^ Rb1^loxP/loxP^* mice have developed histiocytic sarcomas and have not been included in further analyzes. For tamoxifen induction of Cre expression in cell lineage tracing experiments 6 to 8 week-old Slc1a3-CreERT Ai9 mice received 1, 3 and 5 daily i.p. injections of tamoxifen (8 μl/ g body weight, 12.5 mg/ml in corn oil; Sigma-Aldrich, T5648). In tumor induction experiments, 6 to 10 week-old mice Slc1a3-CreERT *Trp53*^loxP/floxP^ *Rb1*^loxP/loxP^ Ai9 mice received i.p. injections every 24 hours for 3 days. K5-CreERT2 *Trp53^loxP/loxP^ Rb1^loxP/loxP^* Ai9 mice of the same age received a single i.p. injection of tamoxifen. All mice were euthanized by CO_2_ and further analyses were carried out.

### Pathological evaluation

All mice in carcinogenesis experiments were subjected to gross pathology evaluation during necropsy. Particular attention was paid to potential sites of ovarian carcinoma spreading, such as the omentum, regional lymph nodes, uterus, liver, lung and mesentery. In addition to the ovary, pathologically altered organs, as well as representative specimens of the brain, lung, liver, kidney, spleen, pancreas and intestine, intra-abdominal lymph nodes, omentum and uterus were fixed in 4% PBS buffered paraformaldehyde. They were then evaluated by microscopic analysis of paraffin sections stained with hematoxylin and eosin and subjected to necessary immunostainings. All early atypical lesions were diagnosed based on their morphology, staining for PAX8 and detection of deleted (floxed-out) *Trp53* and *Rb1* as described earlier^50^. The locations of all lesions were determined by three-dimensional reconstruction of 4-mm-thick serial sections as described previously^51^.

### Immunohistochemistry, apoptosis detection and image analysis

For lineage tracing experiments, Slc1a3-CreERT Ai9 induced mouse tissues were fixed with 4% paraformaldehyde for 1.5 hours on ice. After fixation tissues were washed three times 5 min with PBS at room temperature, embedded in Tissue-Tek O.C.T. compound (ThermoFisherScientific), and frozen on dry ice for 10 min. Immunofluorescent detection of FAM183B and OVGP1 was performed in 7 μm-thick frozen sections according to standard protocols. For paraffin embedding, tissues were fixed in 4% paraformaldehyde overnight at 4°C followed by standard tissue processing, paraffin embedding and preparation of 4 μm-thick tissue sections. For immunohistochemistry, antigen retrieval was performed by incubation of deparaffinized and rehydrated tissue sections in boiling 10 mM sodium citrate buffer (pH 6.0) for 10 minutes. The primary antibodies against PAX8, and tdTomato/RFP were incubated at 4°C for overnight, followed by incubation with secondary biotinylated antibodies (45 minutes, at room temperature, RT). Modified Elite avidin-biotin peroxidase (ABC) technique (Vector Laboratories, Burlingame, CA, USA; pk-6100) was performed at room temperature for 30 minutes. Hematoxylin was used as the counterstain. All primary antibodies used for immunostaining are listed in Supplementary Table 3.

Apoptosis detection was performed on paraffin sections with TUNEL assay following manufacturer instructions (abcam, ab206386), All stained cells were additionally confirmed by morphological recognition of apoptotic bodies, featuring nuclear fragmentation, cytoplasmic shrinkage and blebbing as described earlier ^51,52^. More than 1,200 cells per sample were scored to estimate the apoptotic index.

For quantitative studies, sections were scanned with a ScanScope CS2 (Leica Biosystems, Vista, CA), 40x objective, or Leica TCS SP5 Confocal Microscope, 20x objective, followed by the analysis with Fiji software (National Institutes of Health, Bethesda, MD, USA).

### Microdissection-PCR

Four μm thick paraffin sections were placed on glass slides, stained with H&E and parallel sections stained for tdTomato were evaluated under microdissection microscope. Cells from HE stained sections were collected using a 25G1/2 needle, into 0.6 ml Eppendorf tubes filled with lysis buffer, digested in proteinase K, and processed as described previously ^12,50,51,53^.

### TE organoid preparation

Mouse tubal epithelial cells (TE) were isolated from Slc1a3-CreERT Ai9 positive and negative littermates. Single cell suspensions for organoids were adapted from previously described methods ^7,10,54^. Briefly, mouse uterine tubes were collected, washed with wash buffer (Phosphate buffered saline pH 7.4, Thermo Fisher 10010023, 10,000 U/ml Penicillin-Streptomycin, Thermo Fisher 15140122) and digested with digestion buffer (Gibco DMEM/F12, Fisher 11320033; 4 µg/ml Roche Collagenase/Dispase, Sigma 10269638001; 10 µg/ml DNaseI, Sigma 1128459638001) for two rounds (45 min each) at 37°C. Between rounds, previous digestion buffer was collected and neutralized with 20% FBS containing media in a new tube. New digestion buffer was added to the tissue and mixed vigorously, and digestion allowed to continue. After neutralization, cells were spun down at 600xg for 5 minutes at 4°C and mixed with Matrigel. For every 2 mice (4 uterine tubes), 100 µl of Matrigel would be used and plated along the rim of a 24 well tissue culture plate. Plates were then be incubated in a 37°C cell incubator for 60 minutes, after which Matrigel would be set and organoid media (Gibco Advanced DMEM/F12, Thermo Fisher 12634010; 25% L-WRN Conditioned Medium, ATCC CRL-3276; 12mM HEPES, Thermo Fisher 15630080; 1% GlutaMax, Thermo Fisher 35050079; 2% B27, Thermo Fisher 17504044; 1% N2, Thermo Fisher 17502048; 10 ng/ml hEGF, Sigma E9644; 100 ng/ml Human FGF-10, 1 mM Nicotinamide, Sigma N0636; 10 µM ROCKi, Millipore 688000; 2.5 µM TGF-B RI Kinase Inhibitor VI, Millipore 616464) was added.

For passaging, organoids were released from Matrigel with organoid harvesting media (R&D 3700-100-01) by pipetting around the rim of the plate. Released cells were transferred to a 15 ml falcon tube and allowed to chill on ice for 1 hour. Cells were then spun down in a refrigerated centrifuge at 600xg for 5 minutes at 4°C. Supernatant was aspirated and 1 ml of TrypLE (Thermo Fisher 12604013) was added. Cells were then incubated at 37°C for 10 minutes and vigorously pipetted. Cells were then recovered with 20% FBS containing media and spun down at 600xg for 5 minutes at 4°C. Supernatant was aspirated, organoid media was added, and cells were counted. Typically, cells were plated at 500 cells/µl of Matrigel for expansion. To induce Slc1a3-Cre-ERT mediated recombination, 1 µM 4-Hydroxytamoxifen (Selleck Chem S7827) was supplemented to the media.

### FACS preparation and analysis

Organoid cells were prepared as described above in organoid preparation. After rescuing cells from digestion, cells were washed in 1X PBS + 1% BSA three times. After the last wash, cells were suspended in 1X PBS + 1% BSA. FACS was performed on a Sony MA900 in two separate experiments with single replicates. Sytox blue was used to sort for live cells and Slc1a3-CreERT negative littermates were used as negative controls. Cells were then plated in Matrigel for assessment of organoid forming potential by counting the number of organoids formed against the number of cells sorted. Each replicate contained uterine tubes pooled from at least 5 mice.

### Single-cell RNA-sequencing library preparation

For TE single cell expression and transcriptome analysis we isolated TE from C57BL6 adult estrous female mice. In 3 independent experiments a total of 62 uterine tubes were collected. Each uterine tube was placed in sterile PBS containing 100 IU ml^-1^ of penicillin and 100 µg ml^-1^ streptomycin (Corning, 30-002-Cl), and separated in distal and proximal regions. Tissues from the same region were combined in a 40 µl drop of the same PBS solution, cut open lengthwise, and minced into 1.5-2.5 mm pieces with 25G needles. Minced tissues were transferred with help of a sterile wide bore 200 µl pipette tip into a 1.8 ml cryo vial containing 1.2 ml A-mTE-D1 (300 IU ml^-1^ collagenase IV mixed with 100 IU ml^-1^ hyaluronidase; Stem Cell Technologies, 07912, in DMEM Ham’s F12, Hyclone, SH30023.FS). Tissues were incubated with loose cap for 1 h at 37°C in a 5% CO_2_ incubator. During the incubation tubes were taken out 4 times and tissues suspended with a wide bore 200 µl pipette tip. At the end of incubation, the tissue-cell suspension from each tube was transferred into 1 ml TrypLE (Invitrogen, 12604013) pre-warmed to 37°C, suspended 70 times with a 1000 µl pipette tip, 5 ml A-SM [DMEM Ham’s F12 containing 2% fetal bovine serum (FBS)] were added to the mix, and TE cells were pelleted by centrifugation 300x g for 10 minutes at 25°C. Pellets were then suspended with 1 ml pre-warmed to 37°C A-mTE-D2 (7 mg ml^-1^ Dispase II, Worthington NPRO2, and 10 µg ml^-1^ Deoxyribonuclease I, Stem Cell Technologies, 07900), and mixed 70 times with a 1000 µl pipette tip. 5 ml A-mTE-D2 was added and samples were passed through a 40 µm cell strainer, and pelleted by centrifugation at 300x g for 7 minutes at +4°C. Pellets were suspended in 100 µl microbeads per 10^7^ total cells or fewer, and dead cells were removed with the Dead Cell Removal Kit (Miltenyi Biotec, 130-090-101) according to the manufacturer’s protocol. Pelleted live cell fractions were collected in 1.5 ml low binding centrifuge tubes, kept on ice, and suspended in ice cold 50 µl A-Ri-Buffer (5% FBS, 1% GlutaMAX-I, Invitrogen, 35050-079, 9 µM Y-27632, Millipore, 688000, and 100 IU ml^-1^ penicillin 100 μg ml^-1^ streptomycin in DMEM Ham’s F12). Cell aliquots were stained with trypan blue for live and dead cell calculation. Live cell preparations with a target cell recovery of 5,000-6,000 were loaded on Chromium controller (10X Genomics, Single Cell 3’ v2 chemistry) to perform single cell partitioning and barcoding using the microfluidic platform device. After preparation of barcoded, next-generation sequencing cDNA libraries samples were sequenced on Illumina NextSeq500 System.

### Download and alignment of single-cell RNA sequencing data

For sequence alignment, a custom reference for mm39 was built using the cellranger (v6.1.2, 10x Genomics) *mkref* function. The mm39.fa soft-masked assembly sequence and the mm39.ncbiRefSeq.gtf (release 109) genome annotation last updated 2020-10-27 were used to form the custom reference. The raw sequencing reads were aligned to the custom reference and quantified using the cellranger *count* function.

### Preprocessing and batch correction

All preprocessing and data analysis was conducted in R (v.4.1.1 (2021-08-10)). The cellranger count outs were first modified with the *autoEstCont* and *adjustCounts* functions from SoupX (v.1.6.1) to output a corrected matrix with the ambient RNA signal (soup) removed (https://github.com/constantAmateur/SoupX). To preprocess the corrected matrices, the Seurat (v.4.1.1) *NormalizeData*, *FindVariableFeatures*, *ScaleData*, *RunPCA*, *FindNeighbors*, and *RunUMAP* functions were used to create a Seurat object for each sample (https://github.com/satijalab/seurat). The number of principal components used to construct a shared nearest-neighbor graph were chosen to account for 95% of the total variance. To detect possible doublets, we used the package DoubletFinder (v.2.0.3) with inputs specific to each Seurat object. DoubletFinder creates artificial doublets and calculates the proportion of artificial k nearest neighbors (pANN) for each cell from a merged dataset of the artificial and actual data. To maximize DoubletFinder’s predictive power, mean-variance normalized bimodality coefficient (BC_MVN_) was used to determine the optimal pK value for each dataset. To establish a threshold for pANN values to distinguish between singlets and doublets, the estimated multiplet rates for each sample were calculated by interpolating between the target cell recovery values according to the 10x Chromium user manual. Homotypic doublets were identified using unannotated Seurat clusters in each dataset with the *modelHomotypic* function. After doublets were identified, all distal and proximal samples were merged separately. Cells with greater than 30% mitochondrial genes, cells with fewer than 750 nCount RNA, and cells with fewer than 200 nFeature RNA were removed from the merged datasets. To correct for any batch defects between sample runs, we used the harmony (v.0.1.0) integration method (github.com/immunogenomics/harmony).

### Clustering parameters and annotations

After merging the datasets and batch-correction, the dimensions reflecting 95% of the total variance were input into Seurat’s *FindNeighbors* function with a k.param of 70. Louvain clustering was then conducted using Seurat’s *FindClusters* with a resolution of 0.7. The resulting 19 clusters were annotated based on the expression of canonical genes and the results of differential gene expression (Wilcoxon Rank Sum test) analysis. One cluster expressing lymphatic and epithelial markers was omitted from later analysis as it only contained 2 cells suspected to be doublets. To better understand the epithelial populations, we reclustered 6 epithelial populations and reapplied harmony batch correction. The clustering parameters from *FindNeighbors* was a k.param of 50, and a resolution of 0.7 was used for *FindClusters*. The resulting 9 clusters within the epithelial subset were further annotated using differential expression analysis and canonical markers.

### Pseudotime analysis

Potential of heat diffusion for affinity-based transition embedding (PHATE) is dimensional reduction method to more accurately visualize continual progressions found in biological data ^35^. A modified version of Seurat (v4.1.1) was developed to include the ‘RunPHATE’ function for converting a Seurat Object to a PHATE embedding. This was built on the phateR package (v.1.0.7) (https://github.com/scottgigante/seurat/tree/patch/add-PHATE-again). In addition to PHATE, pseudotime values were calculated with Monocle3 (v.1.2.7), which computes trajectories with an origin set by the user ^36,55–57^. The origin was set to be a progenitor cell state confirmed with lineage tracing experiments.

### Statistical analyses

Statistical comparisons were performed using a two-tailed unpaired *t* test and Analysis of Variance (ANOVA) with Tukey-Kramer Multiple Comparisons Test with InStat 3 and Prism 6 software (GraphPad Software Inc., La Jolla, CA, USA).

## Data Availability

The single-cell RNA-seq data reported in this paper is deposited in Gene Expression Omnibus (GEO); accession number (in process). The Source Data provides data for all results requiring quantification. Any additional data supporting the findings of this study are available from the corresponding author upon reasonable request.

## Code Availability

The fully integrated Seurat objects for the distal and proximal uterine tubes will be available via download on Dryad upon publication. Epithelial subsets of the respective datasets will also be available through the same platform. All code for data preprocessing and figure generation will be made available through GitHub upon publication.

## Acknowledgements

We thank Lora H. Ellenson, Memorial Sloan Kettering Cancer Center, and Matalin Pirtz, Nikitin lab for critical reading of the manuscript, Peter A. Schweitzer, Director of the Cornell Genomics Facility for his invaluable assistance with single-cell RNA sequencing, Tudorita Tumbar, Cornell Molecular Biology and Genetics for providing Slc1a3-CreERT Ai9 mice from her long-term cell lineage tracing experiments and advising on the initial Slc1a3-related experiments, Temirlan Shilikbay, Nazarbayev University, Astana, Kazakhstan for his early contributions to a pilot *Slc1a3* project, and Christopher S. Ashe, Md Mozammal Hossain, Kurtay Ozuner and Derrick Tran for their excellent technical support. This work has been supported by NIH grants (CA182413, CA260115 and CA248524) to AYN, Ovarian Cancer Research Fund grant (327516) to AYN, Sandra Atlas Bass Endowment for Cancer Research to AYN and JCS, and the NSF Graduate Research Fellowship Program (GRFP) awarded to CQR (DGE-2139899).

## Author contributions

AFN and AYN designed experiments. CQR, DJF, DJP, BAH, APA, and SG performed experiments, CQR, DM, and BDC carried our bioinformatics analyses, DJF and AYN performed pathological evaluations, JCS, and BDC provided resources, AFN, CQR and AYN wrote the paper.

## References

1 Siegel, R. L., Miller, K. D., Wagle, N. S. & , Jemal, A. Cancer statistics, 2023. CA Cancer J Clin 73, 17–48 (2023). 10.3322/caac.21763

2 Seidman, J. D., Ronnett, B. M., Shih Ie, M., Cho, K. R. & Kurman, R. J. in Blaustein’s Pathology of the Female Genital Tract (eds R. J. Kurman, L. H. Ellenson, & B. M. Ronnett) 841–966 (Springer, 2019).

3 Carlson, J. W., Gilks, C. B. & Soslow, R. A. Tumors of the ovary and fallopian tube. Vol. 16 (American Registry of Pathology, 2023).

4 Kurman, R. J. & Shih Ie, M. The origin and pathogenesis of epithelial ovarian cancer: a proposed unifying theory. Am J Surg Pathol 34, 433–443 (2010). 10.1097/PAS.0b013e3181cf3d79

5 Landen, C. N., Jr., Birrer, M. J. & Sood, A. K. Early events in the pathogenesis of epithelial ovarian cancer. J Clin Oncol 26, 995–1005 (2008). 10.1200/JCO.2006.07.9970

6 Kim, J. et al. Cell Origins of High-Grade Serous Ovarian Cancer. Cancers (Basel) 10, pii: E433 (2018). 10.3390/cancers10110433

7 Zhang, S. et al. Both fallopian tube and ovarian surface epithelium are cells-of-origin for high-grade serous ovarian carcinoma. Nat Commun 10, 5367 (2019). 10.1038/s41467-019-13116-2

8 Lawrenson, K. et al. A Study of High-Grade Serous Ovarian Cancer Origins Implicates the SOX18 Transcription Factor in Tumor Development. Cell Rep 29, 3726–3735 e3724 (2019). 10.1016/j.celrep.2019.10.122

9 Hao, D. et al. Integrated Analysis Reveals Tubal- and Ovarian-Originated Serous Ovarian Cancer and Predicts Differential Therapeutic Responses. Clin Cancer Res 23, 7400–7411 (2017). 10.1158/1078-0432.CCR-17-0638

10 Lohmussaar, K. et al. Assessing the origin of high-grade serous ovarian cancer using CRISPR-modification of mouse organoids. Nat Commun 11, 2660 (2020). 10.1038/s41467-020-16432-0

11 Maniati, E. et al. Mouse Ovarian Cancer Models Recapitulate the Human Tumor Microenvironment and Patient Response to Treatment. Cell Rep 30, 525–540 e527 (2020). 10.1016/j.celrep.2019.12.034

12 Flesken-Nikitin, A. et al. Ovarian surface epithelium at the junction area contains a cancer-prone stem cell niche. Nature 495, 241–245 (2013). 10.1038/nature11979

13 Yamulla, R. J., Nalubola, S., Flesken-Nikitin, A., Nikitin, A. Y. & Schimenti, J. C. Most Commonly Mutated Genes in High-Grade Serous Ovarian Carcinoma Are Nonessential for Ovarian Surface Epithelial Stem Cell Transformation. Cell Rep 32, 108086 (2020). 10.1016/j.celrep.2020.108086

14 Seidman, J. D. Serous Tubal Intraepithelial Carcinoma Localizes to the Tubal-peritoneal Junction: A Pivotal Clue to the Site of Origin of Extrauterine High-grade Serous Carcinoma (Ovarian Cancer). Int J Gynecol Pathol 34, 112–120 (2015). 10.1097/PGP.0000000000000123

15 Schmoeckel, E. et al. LEF1 is preferentially expressed in the tubal-peritoneal junctions and is a reliable marker of tubal intraepithelial lesions. Mod Pathol 30, 1241–1250 (2017). 10.1038/modpathol.2017.53

16 Kuhn, E. et al. TP53 mutations in serous tubal intraepithelial carcinoma and concurrent pelvic high-grade serous carcinoma--evidence supporting the clonal relationship of the two lesions. J Pathol 226, 421–426 (2012). 10.1002/path.3023

17 Lee, Y. et al. A candidate precursor to serous carcinoma that originates in the distal fallopian tube. J Pathol 211, 26–35 (2007). 10.1002/path.2091

18 Perets, R. et al. Transformation of the fallopian tube secretory epithelium leads to high-grade serous ovarian cancer in Brca;Tp53;Pten models. Cancer Cell 24, 751–765 (2013). 10.1016/j.ccr.2013.10.013

19 Paik, D. Y. et al. Stem-like epithelial cells are concentrated in the distal end of the fallopian tube: a site for injury and serous cancer initiation. Stem Cells 30, 2487–2497 (2012). 10.1002/stem.1207

20 Stewart, C. A. &, Behringer, R. R. Mouse oviduct development. Results Probl Cell Differ 55, 247–262 (2012). 10.1007/978-3-642-30406-4_14

21 Aviles, M., Coy, P. & Rizos, D. The oviduct: A key organ for the success of early reproductive events. Animal Frontiers 5, 25–31 (2015).

22 Ghosh, A., Syed, S. M. & Tanwar, P. S. In vivo genetic cell lineage tracing reveals that oviductal secretory cells self-renew and give rise to ciliated cells. Development 144, 3031–3041 (2017). 10.1242/dev.149989

23 Ford, M. J. et al. Oviduct epithelial cells constitute two developmentally distinct lineages that are spatially separated along the distal-proximal axis. Cell Rep 36, 109677 (2021). 10.1016/j.celrep.2021.109677

24 Flesken-Nikitin, A., Odai-Afotey, A. A. & Nikitin, A. Y. Role of the stem cell niche in the pathogenesis of epithelial ovarian cancers. Mol Cell Oncol 1, e963435 963431–963437 (2014). DOI:10.4161/23723548.2014.963435.

25 Fu, D. J. et al. Stem Cell Pathology. Annu Rev Pathol 13, 71–92 (2018). 10.1146/annurev-pathol-020117-043935

26 Nassar, D. & Blanpain, C. Cancer Stem Cells: Basic Concepts and Therapeutic Implications. Annu Rev Pathol 11, 47–76 (2016). 10.1146/annurev-pathol-012615-044438

27 Fu, D. J. et al. Gastric squamous-columnar junction contains a large pool of cancer-prone immature osteopontin responsive Lgr5(-)CD44(+) cells. Nat Commun 11, 84 (2020). 10.1038/s41467-019-13847-2

28 Friedmann-Morvinski, D. et al. Dedifferentiation of neurons and astrocytes by oncogenes can induce gliomas in mice. Science 338, 1080–1084 (2012). 10.1126/science.1226929

29 Schwitalla, S. et al. Intestinal tumorigenesis initiated by dedifferentiation and acquisition of stem-cell-like properties. Cell 152, 25–38 (2013). 10.1016/j.cell.2012.12.012

30 Rohozinski, J., Diaz-Arrastia, C. & Edwards, C. L. Do some epithelial ovarian cancers originate from a fallopian tube ciliate cell lineage? Med Hypotheses 107, 16–21 (2017). 10.1016/j.mehy.2017.07.014

31 Korsunsky, I. et al. Fast, sensitive and accurate integration of single-cell data with Harmony. Nat Methods 16, 1289–1296 (2019). 10.1038/s41592-019-0619-0

32 Yamanouchi, H., Umezu, T. & Tomooka, Y. Reconstruction of oviduct and demonstration of epithelial fate determination in mice. Biol Reprod 82, 528–533 (2010). 10.1095/biolreprod.109.078329

33 Sada, A. et al. Defining the cellular lineage hierarchy in the interfollicular epidermis of adult skin. Nat Cell Biol 18, 619–631 (2016). 10.1038/ncb3359

34 Ghuwalewala, S. et al. Binary organization of epidermal basal domains highlights robustness to environmental exposure. EMBO J 41, e110488 (2022). 10.15252/embj.2021110488

35 Moon, K. R. et al. Visualizing structure and transitions in high-dimensional biological data. Nat Biotechnol 37, 1482–1492 (2019). 10.1038/s41587-019-0336-3

36 Cao, J. et al. The single-cell transcriptional landscape of mammalian organogenesis. Nature 566, 496–502 (2019). 10.1038/s41586-019-0969-x

37 Xie, Y., Park, E. S., Xiang, D. & Li, Z. Long-term organoid culture reveals enrichment of organoid-forming epithelial cells in the fimbrial portion of mouse fallopian tube. Stem Cell Res 32, 51–60 (2018). 10.1016/j.scr.2018.08.021

38 Rose, I. M. et al. WNT and inflammatory signaling distinguish human Fallopian tube epithelial cell populations. Sci Rep 10, 9837 (2020). 10.1038/s41598-020-66556-y

39 Fu, D. J. et al. Cells expressing PAX8 are the main source of homeostatic regeneration of adult endometrial epithelium and give rise to serous endometrial carcinoma. Dis Model Mech (2020). 10.1242/dmm.047035

40 Network, T. C. G. A. R. Integrated genomic analyses of ovarian carcinoma. Nature 474, 609–615 (2011). 10.1038/nature10166

41 Kaipio, K. et al. ALDH1A1-related stemness in high-grade serous ovarian cancer is a negative prognostic indicator but potentially targetable by EGFR/mTOR-PI3K/aurora kinase inhibitors. The Journal of Pathology 250, 159–169 (2020).

42 Ricciardelli, C. et al. Keratin 5 overexpression is associated with serous ovarian cancer recurrence and chemotherapy resistance. Oncotarget 8, 17819–17832 (2017). 10.18632/oncotarget.14867

43 Logotheti, S. et al. Mechanisms of Functional Pleiotropy of p73 in Cancer and Beyond. Front Cell Dev Biol 9, 737735 (2021). 10.3389/fcell.2021.737735

44 Nemajerova, A. et al. Non-oncogenic roles of TAp73: from multiciliogenesis to metabolism. Cell Death &amp; Differentiation 25, 144–153 (2018). 10.1038/cdd.2017.178

45 Marshall, C. B. et al. p73 Is required for multiciliogenesis and regulates the Foxj1-associated gene network. Cell Reports 14, 2289–2300 (2016). 10.1016/j.celrep.2016.02.035

46 Nemajerova, A. et al. TAp73 is a central transcriptional regulator of airway multiciliogenesis. Genes Dev 30, 1300–1312 (2016). 10.1101/gad.279836.116

47 Song, R. et al. miR-34/449 miRNAs are required for motile ciliogenesis by repressing cp110. Nature 510, 115–120 (2014). 10.1038/nature13413

48 Corney, D. C. et al. Frequent downregulation of miR-34 family in human ovarian cancers. Clin Cancer Res 16, 1119–1128 (2010). 10.1158/1078-0432.CCR-09-2642

49 Levrero, M. et al. The p53/p63/p73 family of transcription factors: overlapping and distinct functions. J Cell Sci 113 **( Pt** **10**), 1661–1670 (2000). 10.1242/jcs.113.10.1661

50 Flesken-Nikitin, A., Choi, K. C., Eng, J. P., Shmidt, E. N. & Nikitin, A. Y. Induction of carcinogenesis by concurrent inactivation of p53 and Rb1 in the mouse ovarian surface epithelium. Cancer Res 63, 3459–3463 (2003).

51 Nikitin, A. Y. & Lee, W. H. Early loss of the retinoblastoma gene is associated with impaired growth inhibitory innervation during melanotroph carcinogenesis in *Rb*^+/-^ mice. Genes & Development 10, 1870–1879 (1996).

52 Nikitin, A. Y. et al. Cell lineage-specific effects associated with multiple deficiencies of tumor susceptibility genes in Msh2(-/-)Rb(+/-) mice. Cancer Res 62, 5134–5138 (2002).

53 Matoso, A., Zhou, Z., Hayama, R., Flesken-Nikitin, A. & Nikitin, A. Y. Cell lineage-specific interactions between Men1 and Rb in neuroendocrine neoplasia. Carcinogenesis 29, 620–628 (2008). https://doi.org/bgm207 [pii] 10.1093/carcin/bgm207

54 Ford, M. J., Harwalkar, K. & Yamanaka, Y. Protocol to generate mouse oviduct epithelial organoids for viral transduction and whole-mount 3D imaging. STAR Protoc 3, 101164 (2022). 10.1016/j.xpro.2022.101164

55 Trapnell, C. et al. The dynamics and regulators of cell fate decisions are revealed by pseudotemporal ordering of single cells. Nature Biotechnology 32, 381–386 (2014). 10.1038/nbt.2859

56 Qiu, X. et al. Single-cell mRNA quantification and differential analysis with Census. Nature Methods 14, 309–315 (2017). 10.1038/nmeth.4150

57 Qiu, X. et al. Reversed graph embedding resolves complex single-cell trajectories. Nat Methods 14, 979–982 (2017). 10.1038/nmeth.4402

