## Supplementary Information for "Pre-ciliated tubal epithelial cells are prone to initiation of high-grade serous ovarian carcinoma"

Flesken-Nikitin et al.

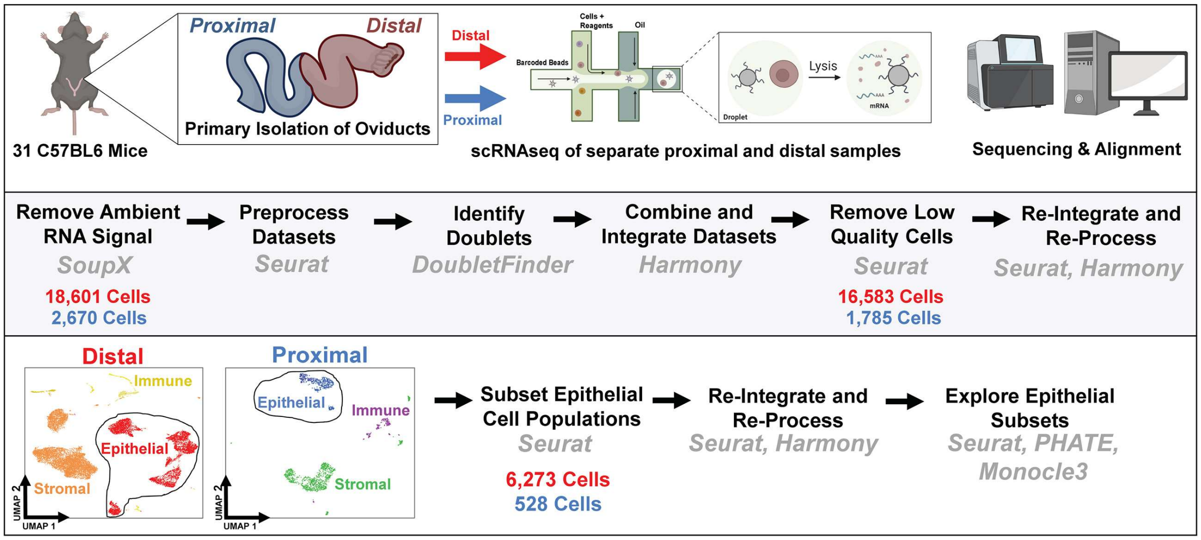

**Supplementary Figure 1. Sample preparation and preprocessing.** After isolation, separation, and disassociation of oviducts to single cell suspensions, the RNA from each cell was sequenced and processed *in silico* to make conclusions of the cell states present within the murine uterine tube.

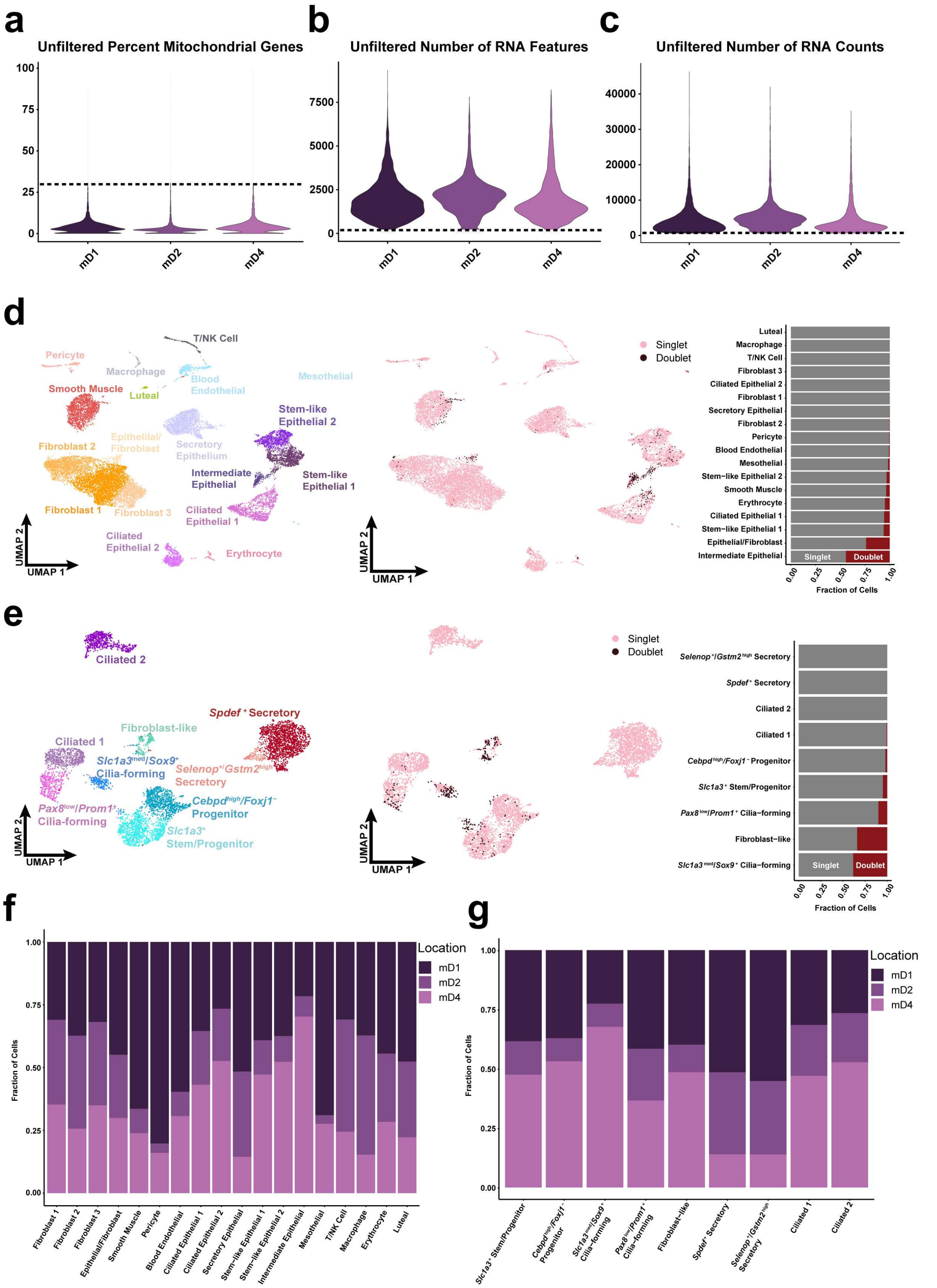

**Supplementary Figure 2. Quality control of scRNA seq datasets.** (**a**) Violin plot demonstrating mitochondrial gene percentage among different sample collection batches with a cutoff of greater than 30%. (**b**) Violin plot demonstrating unique RNA features detected in each cell among different sample collection batches with a cutoff of less than 200. (**c**) Violin plot demonstrating total RNA counts detected in each cell among different sample collection batches with a cutoff of less than 750. (**d**) Doublets detected among cell clusters identified within the complete distal oviduct dataset. (**e**) Doublets detected among cell clusters identified within the epithelial subset. (**f**) Distribution of different sample collection batches attributing to each cluster from the total distal uterine tube dataset. (**g**) Distribution of different sample collection batches attributing to each cluster from the epithelial subset.

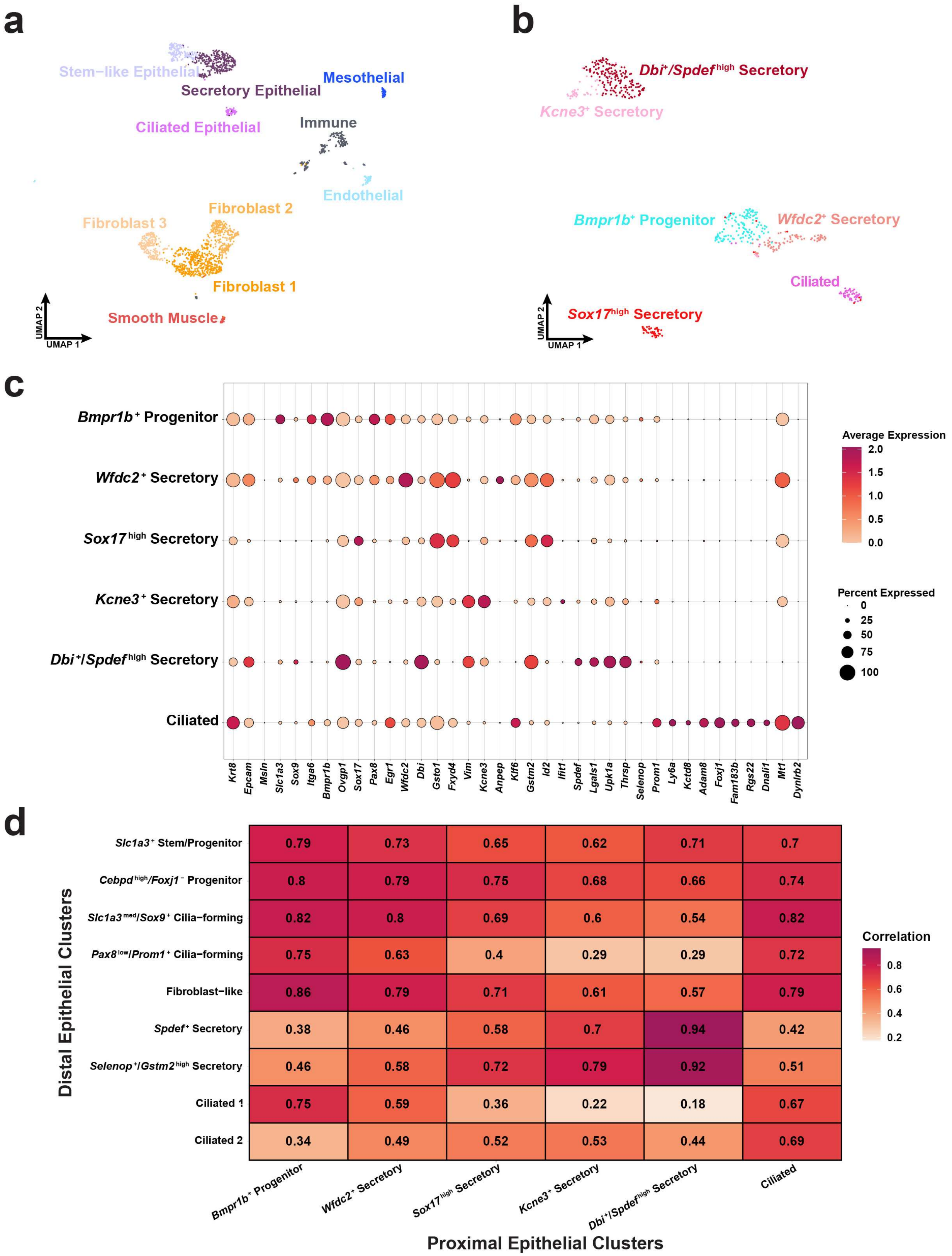

**Supplementary Figure 3. Proximal uterine tube cell atlas.** (**a**) Visualization of 1,785 high quality proximal cells within a UMAP embedding. (**b**) Epithelial cells were identified by their

*Epcam* and *Krt8* expression and a subset of the remaining 528 epithelial cells were represented within the UMAP. (**c**) Dot plot representation of genes associated with the proximal epithelial to validate cell type identification. (**d**) Correlation matrix comparing the average expression profiles of the distal and proximal epithelial clusters.

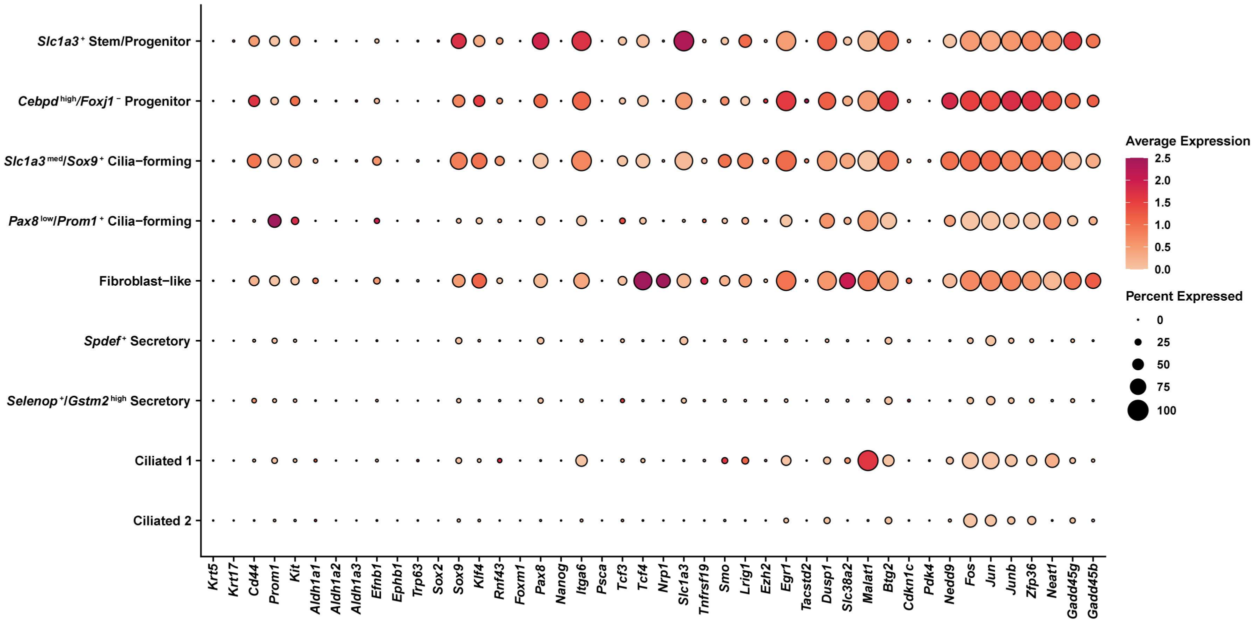

**Supplementary Figure 4. General stem cell marker expression in epithelial cells.** Markers associated with stemness were compiled and used to investigate which epithelial cell clusters may be stem-like.

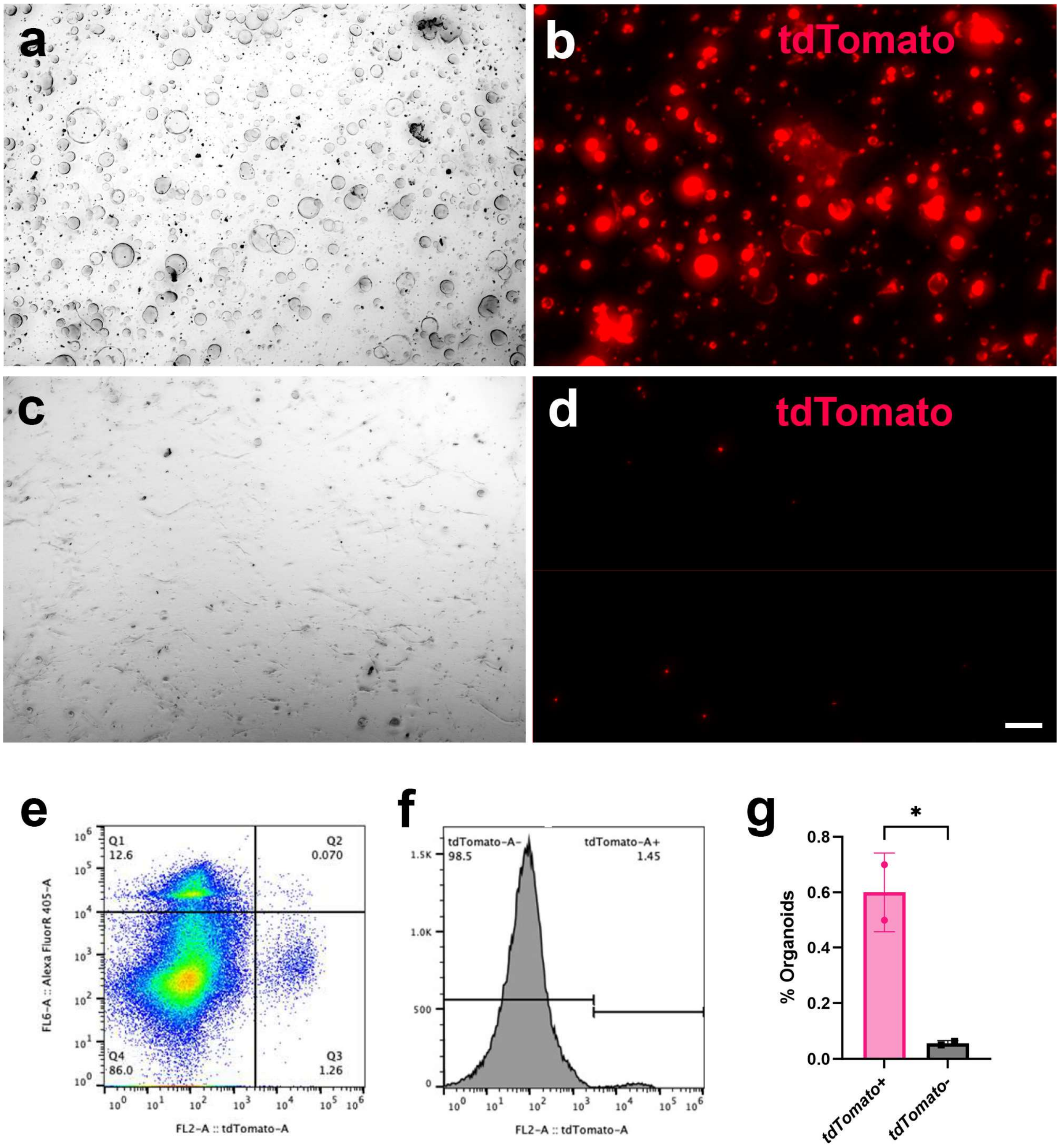

**Supplementary Figure 5. Tubal epithelium (TE) organoid formation.** (**a-d**) Organoid formation by distal (a and b) and proximal (c and d) TE cells isolated from Slc1a3-CreERT Ai9 mice and treated with tamoxifen. Phase contrast (a and c) and tdTomato fluorescence (red, b and d). Bar 100 μm. (**e** and **f**) Representative density plot (e) and histogram (f) of tdTomato+ (Q3, 1.44$\pm$0.688%, mean$\pm$SD, n=4) and tdTomato- (Q4) TE cells separated by FACS 36 hrs after tamoxifen administration. Y-Axis, Sytox Blue. (**g**) Quantification of organoid formation by TE cells FACS separated for tdTomato expression. *unpaired two-tailed P=0.0324 (n=2). Error bar denotes s.d.

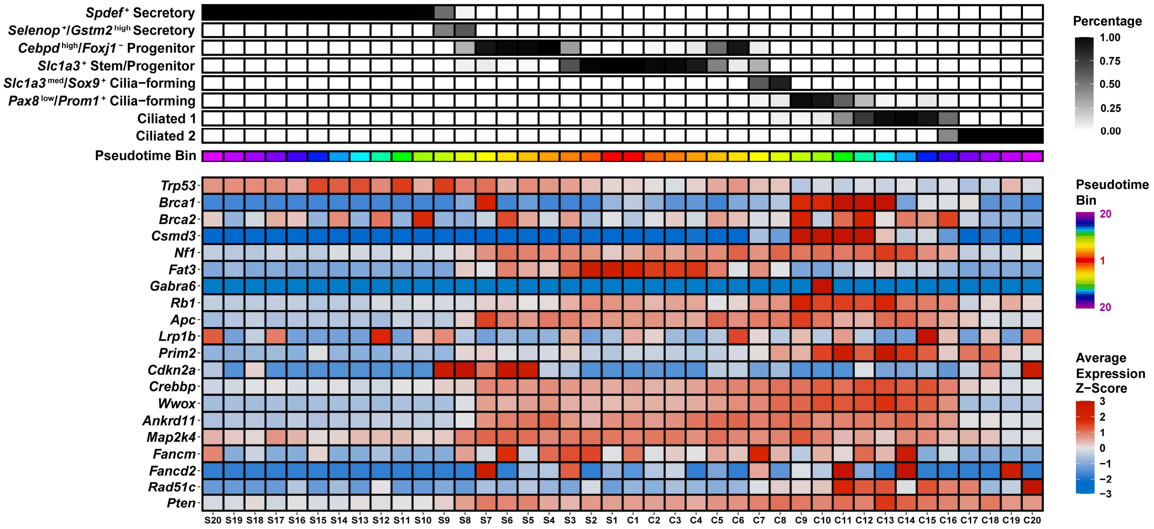

**Supplementary Figure 6. Expression of putative HGSC driver genes along inferred pseudotime trajectories of secretory and ciliated epithelial cell lineages.** The average z-scored expression was calculated for each gene. Each pseudotime bin is equally sized and consists of about 150 cells. Driver genes were derived from the Cancer Genome Atlas Research Network. Most genes, except for FANCM and APC were found to be significantly mutated or deleted in HGSC tumors by TCGA[13, 40]

**Supplementary Table 1.**
Frequency of cell types in the uterine tube

| **Cell States** | **Total Cells** | | **Total *Pax8*+ Cells** | |
|---|---|---|---|---|
| | N | % of Total | N | % of Total |
| **Distal Tubal Epithelium** | | | | |
| Individual Clusters | | | | |
| Ciliated 1 | 714 | 11.38 | 20 | 2.80 |
| Ciliated 2 | 671 | 10.70 | 26 | 3.87 |
| *Pax8*low/*Prom1*+ Cilia-forming | 450 | 7.17 | 168 | 37.33 |
| *Slc1a3*med/*Sox9*+ Cilia-forming | 226 | 3.60 | 163 | 72.12 |
| *Slc1a3*+ Stem/Progenitor | 1126 | 17.95 | 917 | 81.44 |
| *Cebpd*high/*Dbi*+ Progenitor | 935 | 14.91 | 602 | 64.39 |
| *Selenop*+/*Gstm2*high Secretory | 165 | 2.63 | 35 | 21.21 |
| *Spdef*+ Secretory | 1745 | 27.82 | 492 | 28.19 |
| Fibroblast-like | 241 | 3.84 | 158 | 65.56 |
| Total | 6273 | 100 | 2581 | 41.14 |
| Cell Type Groupings | | | | |
| All ciliated and cilia forming cells | 2061 | 32.86 | 377 | 18.29 |
| All secretory cells | 1910 | 30.45 | 527 | 27.59 |
| *Slc1a3*+ Stem/Progenitor cells | 1126 | 17.95 | 917 | 81.44 |
| Transitional cilia-forming cells | 676 | 10.78 | 331 | 48.96 |
| **Proximal Tubal Epithelium** | | | | |
| Individual Clusters | | | | |
| Ciliated | 55 | 10.42 | 16 | 29.09 |
| *Bmp1rb*+ Progenitor | 130 | 24.62 | 88 | 67.69 |
| *Wfdc2*+ Secretory | 80 | 15.15 | 49 | 61.25 |
| *Sox17*high Secretory | 50 | 9.47 | 11 | 22.00 |
| *Kcne3*+ Secretory | 41 | 7.77 | 6 | 14.63 |
| *Dbi*+/*Spdef*high Secretory | 172 | 32.58 | 35 | 20.35 |
| Total | 528 | 100 | 205 | 38.83 |
| Cell Type Groupings | | | | |
| All ciliated and cilia forming cells | 55 | 10.42 | 16 | 29.09 |
| All secretory cells | 343 | 64.96 | 101 | 29.45 |
| Progenitor cells | 130 | 24.62 | 88 | 67.69 |

**Supplementary Table 2.**

Neoplastic TE lesions after Cre-*LoxP* mediated inactivation of *Trp53* and *Rb1*

| Promoter | *Slc1a3* | *Pax8* | *Krt5* |
|---|---|---|---|
| Cases, N | 19 | 21 | 16 |
| First detection (DPI) | NA* | 154 | 104 |
| End point (DPI) | 360 | 400 | 200 |
| Targeted cells (%)# | 11.91±3.23 | 18.89±5.66 | 0.83±0.49 |
| Mice with TE lesions (%) | 0 | 58.33 | 75.00 |

*NA, no applicable, no lesions detected

### According to tdTomato detection in Ai9 mice crossed to "Cre" mice

Mean± s.d., n=6 for *Slc1a3,* n=4 for *Pax8 and Krt5* each. *Pax8 vs Krt5* two-tailed P=0.0007.

**Supplementary Table 3.**

List of antibodies used for immunostaining

| Antigen | Antibody source, catalogue number | Clone | Host | Dilution | Detection system |
|---|---|---|---|---|---|
| FAM183B | Invitrogen, PA5-71109 | PC* | Rabbit | 1:100 | Immunofluorescence |
| PAX8 | Protein Tech., 10366-1-AP | PC | Rabbit | 1:2000<br>1:400 | Vectastain Elite ABC-HRP Kit<br>Immunofluorescence |
| RFP | Rockland Immunochemicals Inc., 600-401-379S | PC | Rabbit | 1:400<br>1:100 | Vectastain Elite ABC-HRP Kit<br>Immunofluorescence |
| OVGP1 | Abcam, Ab118590 | PC | Rabbit | 1:500 | Immunofluorescence |
| Anti-rabbit IgG (Biotinylated) | Vector Laboratories, BA-1000 | PC | Goat | 1:200 | Vectastain Elite ABC-HRP Kit |
| Anti-rabbit IgG conjugated with Alexa 488 | Invitrogen, A21206 | PC | Donkey | 1:200 | Immunofluorescence |
| Anti-rabbit IgG conjugated with Alexa 594 | Invitrogen, A21207 | PC | Donkey | 1:200 | Immunofluorescence |

*PC: Polyclonal
